# HDAC4-mediated deacetylation of GSDMD prevents pyroptosis by inhibiting GSDMD ubiquitination

**DOI:** 10.1101/2023.06.02.543413

**Authors:** Weilv Xu, Xinyue Li, Danyue Li, Xinyu Fu, Nan Chen, Qian Lv, Yuhua Shi, Suhui He, Lu Dong, Yang Yang, Fushan Shi

## Abstract

Gasdermin D (GSDMD) functions as a key pyroptotic executor and induces cytokine secretion after cleavage by inflammatory caspases. However, less is known about the role of posttranslational modifications (PTMs) in GSDMD-mediated pyroptosis. Here, we report that GSDMD can be acetylated at Lysine 248 residue and the acetylation of GSDMD promotes pyroptosis. We identified histone deacetylase 4 (HDAC4) as the specific deacetylase that mediates GSDMD deacetylation and subsequent pyroptosis inhibition *in vitro* and *in vivo*. GSDMD deacetylation impairs the ubiquitination of GSDMD, for which pyroptosis is inhibited. Interestingly, the phosphorylation of HDAC4 is important for its ability of deacetylating GSDMD and suppressing GSDMD-mediated pyroptosis. The Protein phosphatase 1 (PP1) catalytic subunits (PP1α and PP1γ) mediate the dephosphorylation of HDAC4, thereby abrogating its deacetylase activity on GSDMD. Therefore, our work unravels a complex regulatory mechanism involving HDAC4, PP1 and GSDMD, and provides novel insights into the crosstalk among acetylation, ubiquitination and phosphorylation.

## Introduction

Pyroptosis is a type of programmed necrotic cell death, featuring cell swelling, and large bubbles blowing from the plasma membrane, and has received increasing attention for its association with innate immunity^1-3^. Pyroptosis is mediated by inflammasomes assembly and activation^4,5^. Canonical inflammasomes, including NLRP1, NLRP3, NLRC4 and AIM2 inflammasome, are assembled in the cytosol to recruit and activate caspase-1. Noncanonical inflammasomes are activated by human caspase-4/5 (mouse orthologs caspase-11) directly binding to intracellular lipopolysaccharide (LPS)^5-8^.

Gasdermins are reported to be the executioner of pyroptosis. The gasdermin family is composed of Gasdermin A-E and DFNB59, which shares the propyroptotic activity and autoinhibited structure^1-3,9^. Among them, Gasdermin D is most widely-investigated. GSDMD could be cleaved by caspase-1/4/5/11 and liberates its N-terminal domain (GSDMD-p30), which can oligomerize to form pores in the cell membrane, causing the release of inflammatory cytokines and triggering pyroptosis^10-13^.

Recently, posttranslational modifications (PTMs) have been shown to play a significant role in the regulation of pyroptosis. Many components of inflammasomes are reported to be affected by posttranslational modifications. SIRT2-mediated deacetylation of NLRP3 could inactivate the NLRP3 inflammasome while SIRT3 deacetylates NLRC4 to promote NLRC4 inflammasome activation^14,15^. Moreover, studies have shown that the function of GSDMD on pyroptosis could be modified by ubiquitination. Our previous work shows that SYVN1 mediates K27-linked polyubiquitination of GSDMD to promote the pyroptotic cell death^16^. Shigella ubiquitin ligase IpaH7.8 promotes GSDMD degradation via ubiquitination to prevent pyroptosis^17^. In addition, there is a connection between posttranslational modifications. Acetylation of OTUD3 significantly enhances its activity for the Lys63 linkage on MAVS and ubiquitination of p62 can be promoted by acetylation^18,19^. So far, there has been little research directly investigating the acetylation of GSDMD. Furthermore, the relationship between posttranslational modifications of GSDMD is also needed to be elucidated.

In this study, we report that GSDMD could be acetylated and histone deacetylase 4 (HDAC4) inhibits pyroptosis through deacetylating GSDMD. Moreover, we identify the Protein phosphatase 1 (PP1) as the phosphatase for HDAC4. The phosphorylation of HDAC4 shows significant effects on its activity in suppressing pyroptosis. Intriguingly, the acetylation of GSDMD promotes its ubiquitination, a mechanism not yet described for pyroptosis.

## Results

### Acetylation of GSDMD promotes pyroptosis

Previous studies have shown that acetylation plays significant roles in the regulation of inflammation^15,20-22^. Given that GSDMD is a key downstream effector of inflammasomes, we intend to find whether GSDMD undergoes acetylation/deacetylation during pyroptosis. In order to confirm whether GSDMD could be acetylated, we analyzed the acetylation of exogenously expressed GSDMD in HEK293T cells treated with trichostatin A (TSA), a broad-spectrum inhibitor of HDAC family deacetylases, and nicotinamide (NAM), an inhibitor of SIRT family deacetylases. By performing immunoprecipitation, using a specific antibody against acetylated lysine, we detected strong acetylation of GSDMD in TSA-treated but not NAM-treated cells (Fig. 1a). Additionally, in THP-1 cells treated with LPS and nigericin, which activated the canonical NLRP3 inflammasome, we found the acetylation of endogenous GSDMD was evidently increased (Fig. 1b, c). We then tested whether GSDMD from other species could be acetylated or not. We confirmed the acetylation of GSDMD from the pig and the mouse in IPEC-J2 cells and RAW264.7 cells (Fig. 1d, e). HEK293T cells transfected with p-GSDMD and m-GSDMD were also tested (Fig. 1f). These results suggest that GSDMD is an acetylated protein.

**Fig. 1.**
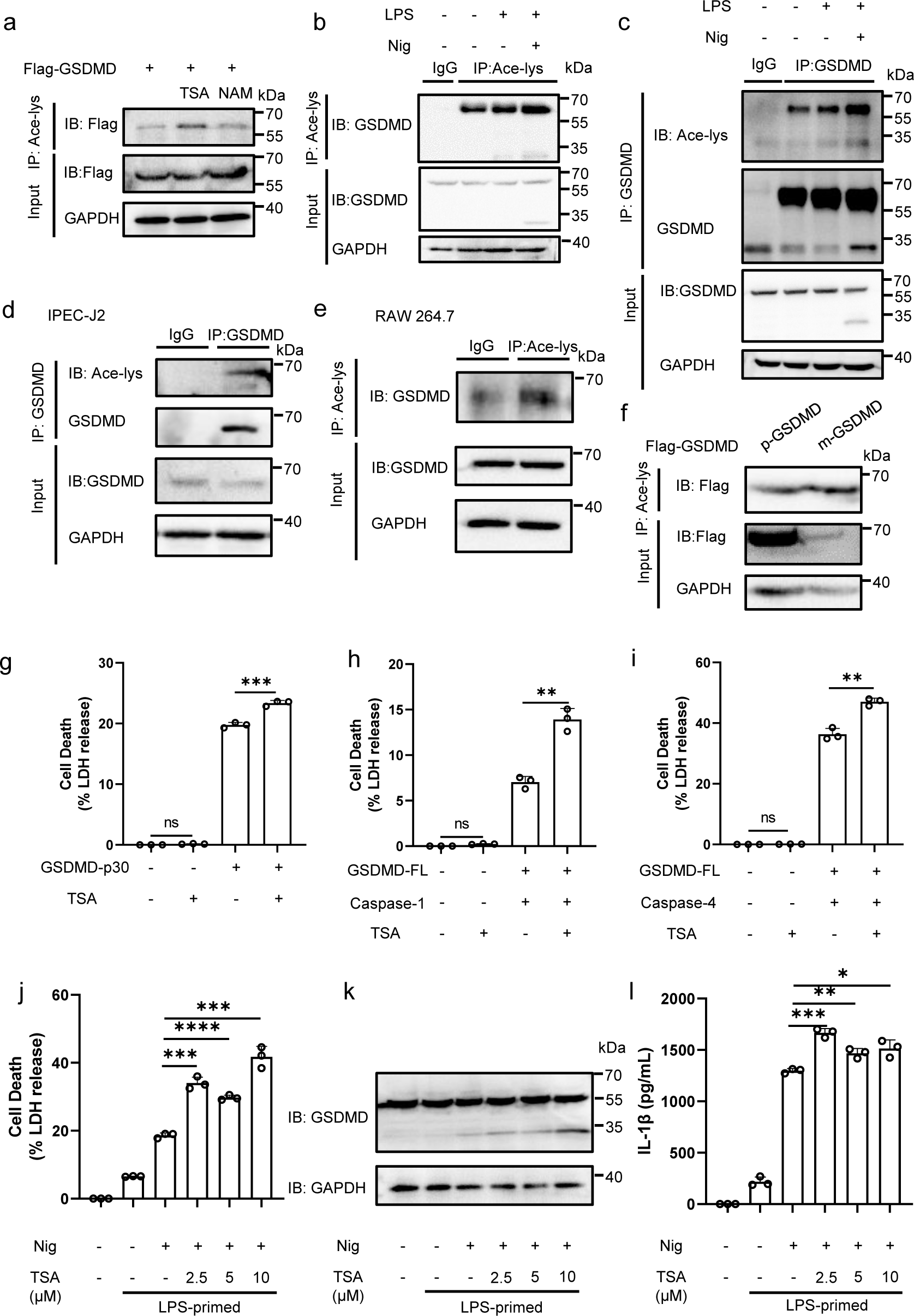
Acetylation of GSDMD affects pyroptosis. a. Acetylation of exogenous Flag-GSDMD in HEK293T cells treated with deacetylase inhibitors TSA or NAM. Flag-GSDMD was immunoprecipitated with anti-acetyl-lys antibody (Ace-lys), and the precipitates were analyzed using an anti-flag antibody. **b-e** Acetylation of endogenous GSDMD in THP-1 (**b, c**), IPEC-J2 (**d**), RAW264.7 (**e**) cells. GSDMD acetylation was analyzed by immunoprecipitation with an anti-acetyl-lys antibody (**b, d, e**) or anti-GSDMD antibody (**c**) followed by western blotting. **f** Acetylation of exogenous mouse-Flag-GSDMD and pig-Flag-GSDMD in HEK293T cells. **g-i** HEK293T cells transfected with GSDMD-p30 or GSDMD-FL (full length) and Caspase-1 or GSDMD-FL and Caspase-4 were treated with TSA (10 μM). LDH release was analyzed using an LDH release assay. **j-l** THP-1 cells were incubated with 500 ng/mL LPS and TSA for 4 h and then another 1 h for 10 μM Nigericin. The supernatants were collected and analyzed by LDH release assay (**j**) and ELISA for IL-1β (**l**). Cell lysates were analyzed by immunoblotting (**k**). ****stands for P<0.0001, ***stands for P<0.001, ** stands for P<0.01, * stands for P<0.05 and ns stands for no significant difference (unpaired t test). Data shown are mean ± SD from one representative experiments performed in triplicate.

As GSDMD is an executor of pyroptosis, we next evaluated the effect of GSDMD acetylation on pyroptosis. HEK293T cells were respectively transfected with GSDMD-p30 or GSDMD-full length (GSDMD-FL) and Caspase-1 or GSDMD-FL and Caspase-4, treated with or without TSA. We found that TSA treatment could remarkably increase the release of LDH and propidium iodide (PI) staining, suggesting that pyroptosis was promoted (Fig. 1g-i and Supplementary Fig.1). HEK293T cells transfected with p-GSDMD or m-GSDMD obtained similar results (Supplementary Fig. 2a-d). In addition, pyroptotic cell death triggered by inflammasome activation was also checked. We first treated THP-1 cells with TSA to avoid the interference of drug toxicity on the experiment (Supplementary Fig. 3a, b). THP-1 cells were primed with LPS and treated with NLRP3 inflammasome activator Nigericin (Nig), and then LDH release, expression of GSDMD and IL-1β secretion were detected. LPS and Nig treatment induced cell death and cleavage of GSDMD, which could be strongly promoted by TSA-treatment (Fig. 1j-l). Moreover, THP-1 primed with Pam3CSK4 and followed by LPS transfection to trigger noncanonical NLRP3 inflammasome showed similar results (Supplementary Fig. 2e-g). To examine the effect of TSA on other inflammasomes, we utilized poly (dA:dT), a trigger of the AIM2 inflammasome, and flagellin, an activator of the NLRC4 inflammasome. As shown in Supplementary Fig. 2h-m, we found that the release of LDH, cleavage of GSDMD and the secretion of IL-1β triggered by inflammasome activators in THP-1 cells treated with TSA were significantly increased. Thus, these results suggest that the acetylation of GSDMD promotes pyroptosis.

### Acetylation of the K248 residue of GSDMD modulates pyroptosis

To find the potential acetylation sites, we used mass spectrometry analysis of Flag-GSDMD from TSA-treated HEK293T cells and GPS-PAIL 2.0^23^, a software to predict lysine modification sites. The results revealed four potential lysine residues: K103, K145, K248 and K387. Sequence comparison revealed that Lys103, Lys145 and Lys248 are conserved acetylation motif in GSDMD orthologs (Supplementary Fig. 4). Thus, we constructed several mutants (K103R, K145R, K248R and K387R) based on the GSDMD-full length or GSDMD-p30. As shown in Fig. 2a-c, K103R and K248R mutants showed decreased pyroptotic activity than wildtype GSDMD (GSDMD-WT), while K145R and K387R mutants not. Compared to GSDMD-WT, only GSDMD-K248R mutant showed markedly reduced acetylation (Fig. 2d), suggesting Lysine 248 residue was one of the key acetylation sites of GSDMD.

**Fig. 2.**
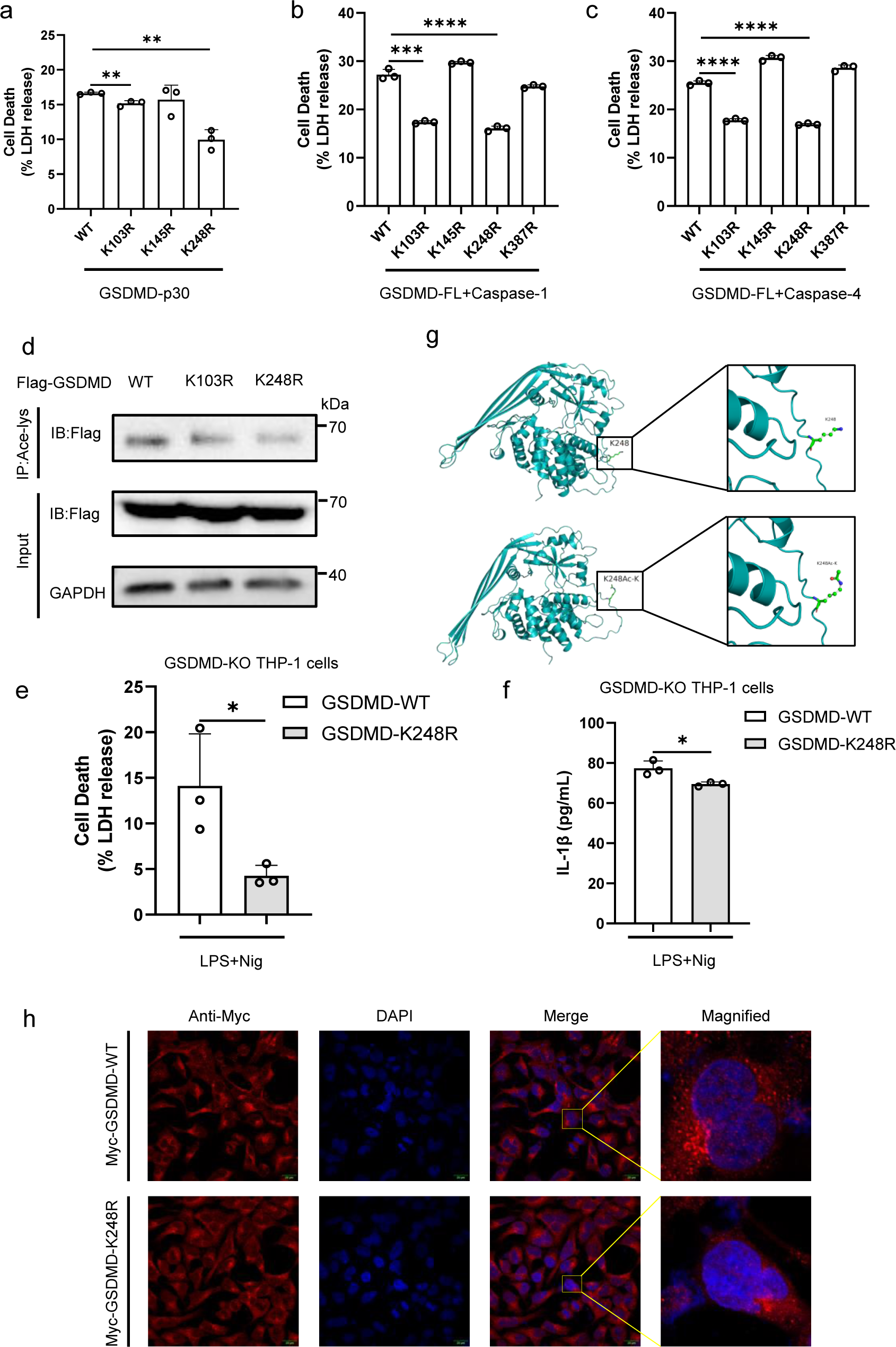
GSDMD is acetylated at Lysine 248. **a-c** HEK293T cells were transfected with GSDMD-p30 mutants (**a**) or GSDMD-FL mutants and Caspase-1 (**b**) or GSDMD-FL mutants and Caspase-4 (**c**). The release of LDH in the culture medium was detected. **d** Acetylation of GSDMD mutants expressed in HEK293T cells. **e-f** PMA-differentiated GSDMD-KO THP-1 macrophages were infected with Flag-GSDMD-FL-WT/K248R-Lentivirus, followed with LPS and nigericin treatment. The supernatants were collected and analyzed by LDH release assay (**e**) and ELISA for IL-1β (**f**). **g** Domain organization and ribbon diagrams of GSDMD-WT (left) and GSDMD-K248Ac-K (right). The acetylated GSDMD structure was constructed using the Structure Editing module of UCSF Chimera and optimized using Minimize Structure. **h** Immunofluorescence microscopy and nuclear staining (with the DNA-binding dye DAPI) of HEK293T cells transfected with expression plasmids for Myc-GSDMD-WT or Myc-GSDMD-K248R. Scale bars, 20 μm. ****stands for P<0.0001, ***stands for P<0.001, ** stands for P<0.01, * stands for P<0.05 (unpaired t test). Data shown are mean ± SD from one representative experiments performed in triplicate.

Additionally, reconstruction of GSDMD-K248R in GSDMD-KO THP-1 cells caused milder cell death than GSDMD-WT (Fig. 2e, f). Through structural modeling, we observed acetyl group could deeply dock with GSDMD at K248 residue (Fig. 2g). Immunofluorescence showed that GSDMD-K248R did not affect its localization compared to GSDMD-WT (Fig. 2h). Therefore, these results indicate that GSDMD is acetylated at Lys 248.

### Histone deacetylase 4 inhibits GSDMD-mediated pyroptosis

As mentioned above, the acetylation of GSDMD was evidently increased by the stimulation of TSA rather than NAM, suggesting the involvement of HDAC family deacetylases. To identify the specific deacetylase of GSDMD, we performed a mass spectrometry analysis of Flag-tagged GSDMD with or without TSA treatment. This analysis identified histone deacetylase 4 (HDAC4) as a specific deacetylase of GSDMD (Fig. 3a). Co-immunoprecipitation analysis showed a significant interaction between GSDMD and HDAC4 in HEK293T cells (Fig. 3b). Interaction of endogenous HDAC4 with GSDMD was also confirmed in THP-1 cells (Fig. 3c). This interaction was also observed in pigs and mice (Supplementary Fig. 5a, b). Immunofluorescence analysis showed that GSDMD and HDAC4 co-localized well in cytoplasm (Supplementary Fig. 5c). We then found HDAC4 could significantly inhibit pyroptosis induced by GSDMD derived from different species (Fig. 3f and Supplementary Fig. 5d-i). HDAC4 co-transfection also reduced PI intake (Supplementary Fig. 6). Moreover, we transfected HEK293T cells with sg-HDAC4 and tested its efficiency (Supplementary Fig. 5j). Then we infected THP-1 cells with the viral supernatant from HEK293T cells. We found that sg-HDAC4 THP-1 cells showed severer cell death after stimulation of LPS and Nig (Fig. 3d, e and Supplementary Fig. 5k). To further certify the above findings, an *in vivo* model was employed. First, the efficiency of HDAC4 knock-down by siRNA was verified at the protein level in NIH-3T3 cells (Supplementary Fig. 5l). Then si-HDAC4#1 or negative control siRNA was intraperitoneally injected into mice, and the expression of HDAC4 decreased in various organs (Fig. 3g). Mice were then injected with LPS (20 mg/kg) for 6 h before serum and peritoneal lavage fluid were collected. We found that the knock-down of HDAC4 inhibited the secretion of IL-1β, while the production of TNF-α and IL-6 was not affected (Fig. 3h-m).

**Fig. 3.**
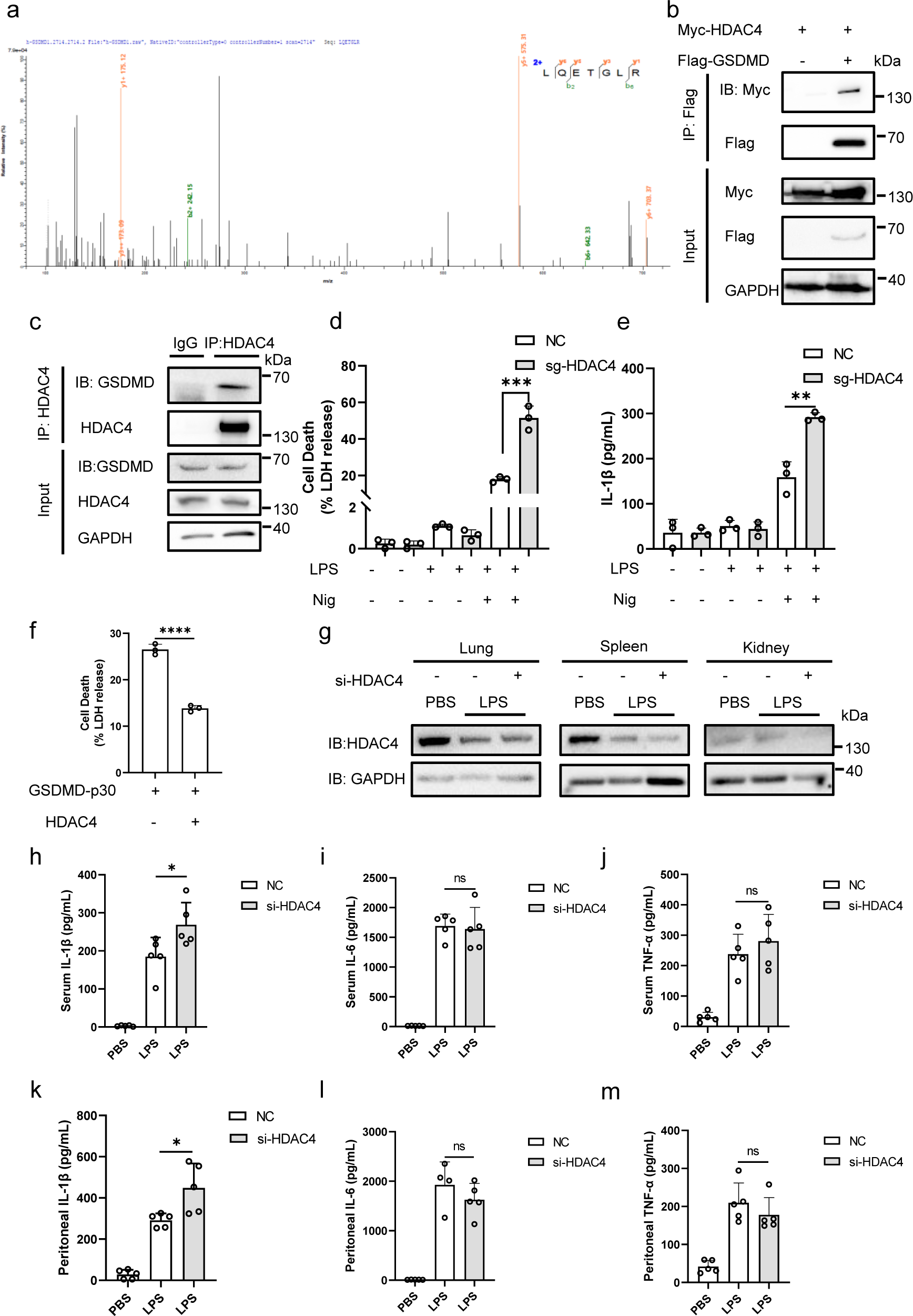
HDAC4 inhibits pyroptosis. **a** Mass spectrometry analysis of Flag-tagged GSDMD immunoprecipitated by anti-Flag antibody in HEK293T cells treated with TSA. **b** IB of total cell lysates (input) and proteins immunoprecipitated with anti-Flag resin from HEK293T cells transfected with Myc-HDAC4 and Flag-GSDMD. **c** IB of total lysates (input) and proteins immunoprecipitated with control or anti-HDAC4 antibodies in THP-1 cells. **d-e** LDH release (**d**) and ELISA (**e**) of IL-1β in NC or sg-HDAC4 THP-1 cells treated with LPS and Nig. **f** LDH release of HEK293T cells transfected with GSDMD-p30, together with or without HDAC4. **g** Immunoblot analysis of the protein level of HDAC4 in various organs of control or HDAC4-knockdown mice after LPS injection for 6 h. **h-m** ELISA analysis of IL-1β, TNF-α and IL-6 secretion from the serum or peritoneal lavage fluid of control or HDAC4-knockdown mice after LPS injection for 6 h. (mice: n = 5 for each group). ****stands for P<0.0001, * stands for P<0.05 and ns stands for no significant difference (unpaired t test). Data shown are mean ± SD from one representative experiments performed in triplicate.

Moreover, we used LMK-235, an inhibitor of HDAC4, to confirm the participation of HDAC4 in deacetylation of GSDMD. The treatment of LMK-235 could promote LDH release in HEK293T cells transfected with GSDMD-p30 or GSDMD-FL and Caspase-1 or GSDMD-FL and Caspase-4 (Supplementary Fig. 7a-c). We also treated THP-1 cells with LMK-235 to check if the drug had effect on the release of LDH and IL-1β (Supplementary Fig. 3c, d). As shown in Supplementary Fig. 7d-o, LMK-235 showed similar effects on inflammasomes as TSA, which supports the observation that HDAC4 prevents NLRP3, AIM2 and NLRC4 inflammasome-mediated pyroptosis. The results above suggest that HDAC4 inhibits pyroptosis both *in vivo* and *in vitro*.

### HDAC4 interacts with GSDMD in multiple domains

Though we had demonstrated the interaction between HDAC4 and GSDMD (Fig. 3b, c), we still intended to determine which domain of HDAC4 is critical for its interaction with GSDMD. Then we created different deletion mutants of HDAC4 (Fig. 4a) and examined their co-immunoprecipitation with GSDMD. We found that all the mutants could bind with GSDMD, suggesting there were several binding sites between GSDMD and HDAC4 (Fig. 4b). Using structural modeling, we analyzed the binding of GSDMD K248Ac-K with HDAC4 (Fig. 4c, d). The analysis showed that there are multiple interactions between them. As shown in Fig. 4d, the R249 of GSDMD and the negative-charged D354 on HDAC4 formed a salt bridge interaction; negative-charged D228 of GSDMD and the positive-charged R51 on HDAC4 formed a salt bridge and hydrogen bond interaction. Q237 and S225 of GSDMD formed hydrogen bonds with Q46 and C54 on HDAC4. The acetylated K248 also has van der Waals contact with N351 and E352 on HDAC4, thereby facilitating substrate bindings. Then we investigated the effects of those HDAC4 mutants on pyroptosis. HEK293T cells were transfected with GSDMD-p30 or GSDMD-FL and Caspase-1 or GSDMD-FL and Caspase-4, together with HDAC4-FL or deletion mutants. The release of LDH showed that HDAC4 remarkably inhibited pyroptosis while mutants could rescue the suppression of HDAC4 to some extent, which also supports that HDAC4 binds to GSDMD in multiple sites (Fig. 4e-g).

**Fig. 4.**
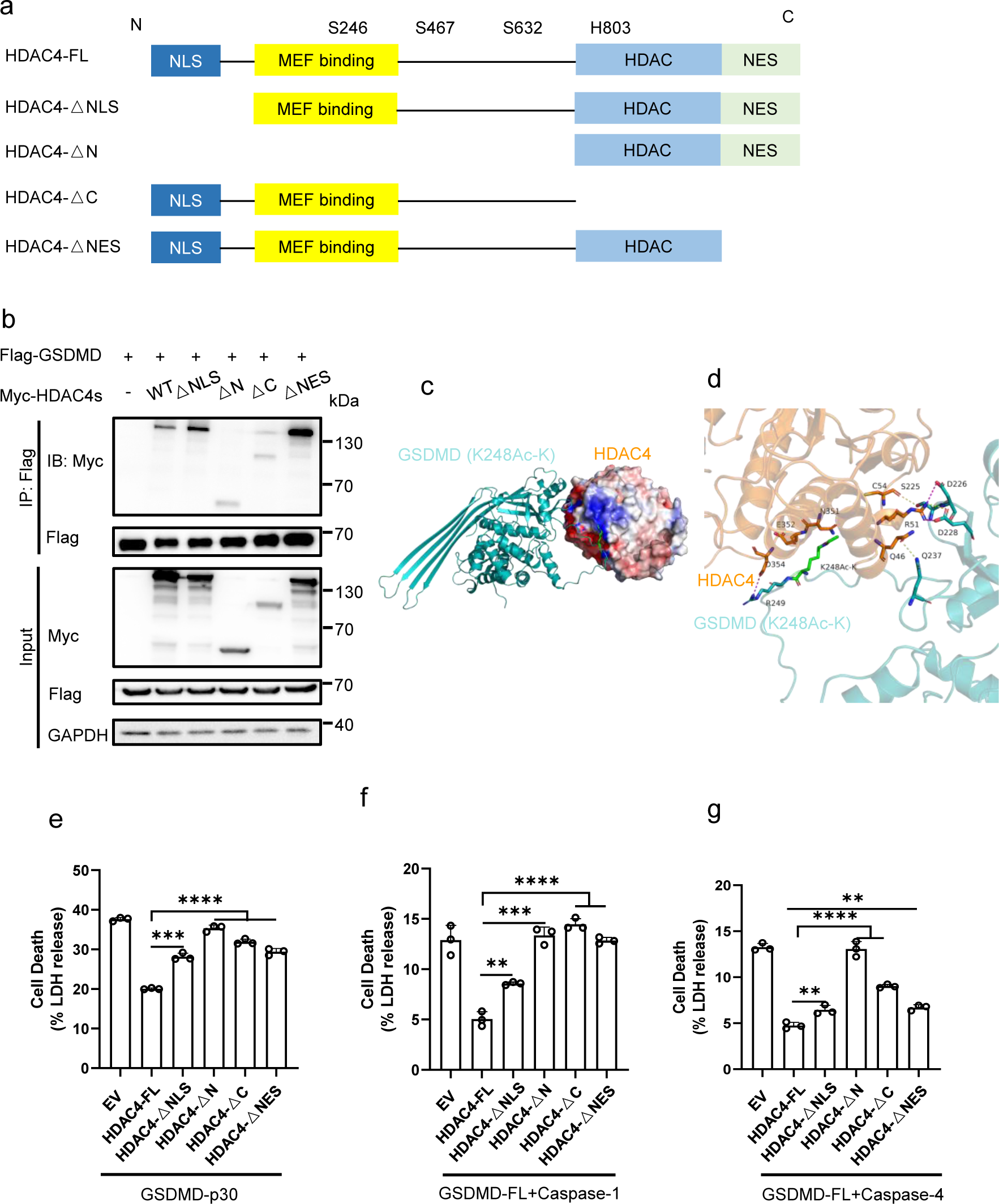
Mapping the functional domain of HDAC4 in GSDMD-mediated pyroptosis. **a** Schematic representation of HDAC4 and its mutants. **b** IB of total cell lysates (input) and proteins immunoprecipitated with anti-Flag resin from HEK293T cells transfected with Myc-HDAC4 or its mutants and Flag-GSDMD. **c-d** Structural model of the interaction of K248-acetylated GSDMD and HDAC4 (PDB: 2VQW). Blue represents the electropositive region, and red represents electronegative region (**c**). 3D display of interaction key amino acids (**d**), hydrogen bonds are represented by yellow dotted lines, and salt bridges are represented by magenta dotted lines. Acetylated lysine K248Ac-K was highlighted with green. The docking models were analyzed using HDOCK Server. **e-g** LDH release of HEK293T cells transfected with GSDMD-p30 (**e**) or GSDMD-FL and Caspase-1 (**f**) or GSDMD-FL and Caspase-4 (**g**), together with HDAC4-WT or its mutants. ****stands for P<0.0001, ***stands for P<0.001, ** stands for P<0.01 (unpaired t test). Data shown are mean ± SD from one representative experiments performed in triplicate.

### GSDMD is deacetylated by HDAC4

We next determined whether the inhibitory effect of HDAC4 on pyroptosis depended on its enzymatic activity. We overexpressed HDAC4 in HEK293T cells and found that the acetylation of GSDMD was greatly reduced, which could be totally rescued by TSA (Fig. 5a). In contrast, the acetylation level of GSDMD in sg-HDAC4 HEK293T cells increased evidently (Fig. 5b). Moreover, plasmid encoding HDAC4-H803L (abolishing HDAC enzymatic activity) could not deacetylate GSDMD (Fig. 5c). However, we detected that HDAC4-H803L could not inhibit pyroptosis (Fig. 5d-f). As HDAC4-H803L could not deacetylate GSDMD but still could inhibit pyroptosis, we hypothesized that other sites on HDAC4 played key roles in inhibition of pyroptosis. Since HDAC4 shuttles between nucleus and cytoplasm^24^, and the phosphorylation of HDAC4 at Ser 246, Ser 467 and Ser 632 is crucial for it to export to cytoplasm^25,26^. We then constructed HDAC4-S246A, S467A, S632A, and S246/467/632A (3SA) mutants. We found that these three sites all had effects on pyroptosis and the 3SA mutant almost completely lost the inhibitory capacity on pyroptosis (Fig. 5d-f), while the phosphomimetic mutant (3SE) was still capable of inhibiting pyroptosis (Fig. 5h-j). As we found in Supplementary Fig. 5c, HDAC4 was distributed within the cytoplasm and the nucleus, while the co-expression of HDAC4 and GSDMD led to a redistribution of HDAC4 to the cytoplasm. It is possible that phosphomimetic mutation of HDAC4 abrogated its transfer from nucleus to cytoplasm, which reduced the opportunity for HDAC4 and GSDMD interaction. Furthermore, we observed that the three residues were also important for the deacetylation activity of HDAC4 (Fig. 5g). Thus, our data implies that S246, S467 and S632 residues of HDAC4 were functionally important for HDAC4-mediated deacetylation of GSDMD and inhibition of pyroptosis.

**Fig. 5.**
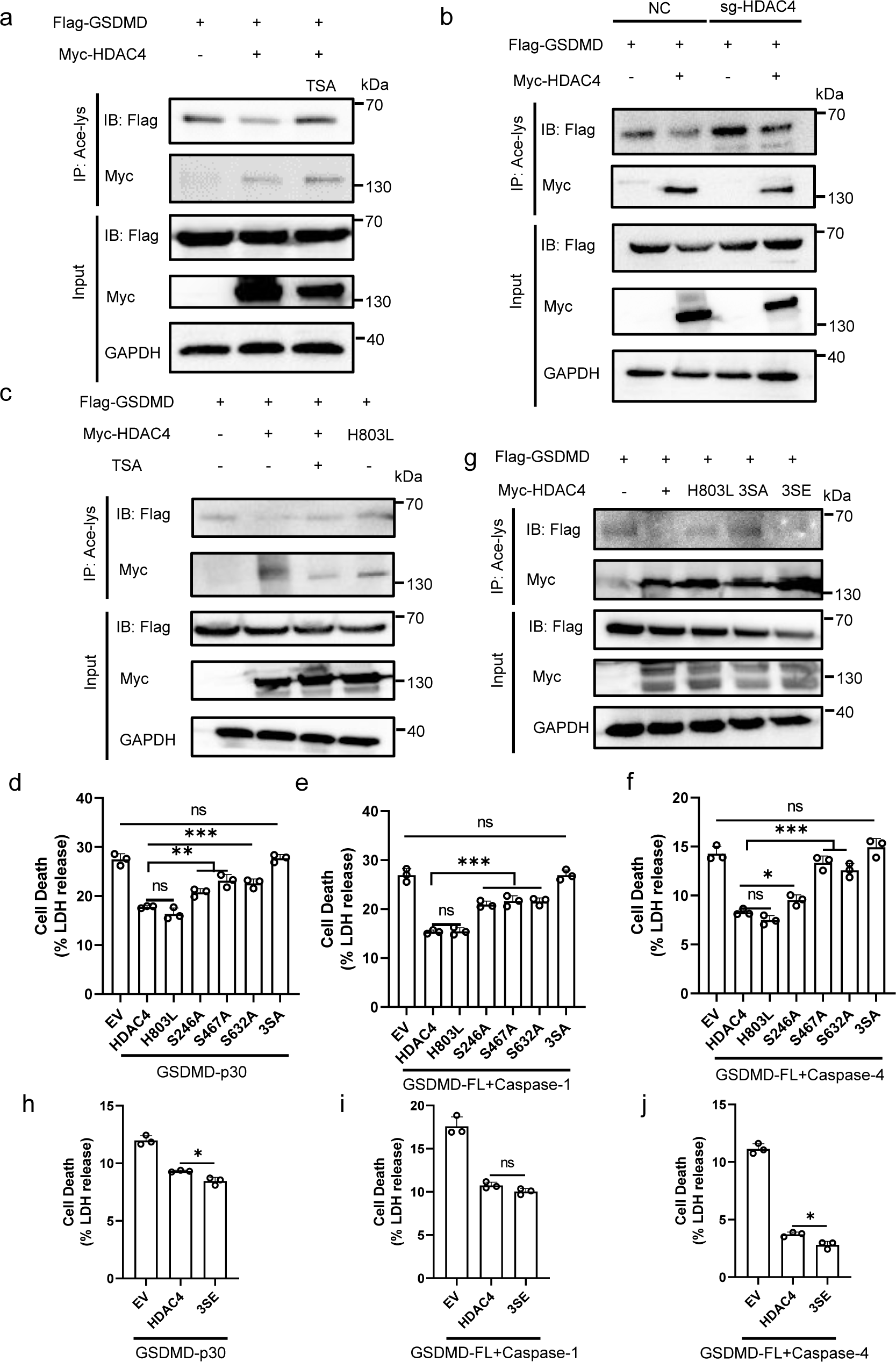
HDAC4 deacetylates GSDMD. **a** HEK293T cells were transfected with Flag-GSDMD and Myc-HDAC4, treat with or without TSA. Proteins were immunoprecipitated with anti-acetyl-lys antibody (Ace-lys), and the precipitates were analyzed using an anti-Flag or anti-Myc antibody. **b** NC and sg-HDAC4 HEK293T cells were transfected with Flag-GSDMD, together with or without Myc-HDAC4. GSDMD acetylation was analyzed by immunoprecipitation with an anti-acetyl-lys antibody followed by western blotting. **c** HEK293T cells were transfected with Flag-GSDMD and Myc-HDAC4 or HDAC4-803L, treated with or without TSA. GSDMD acetylation was analyzed by immunoprecipitation. **d-f** LDH release of HEK293T cells transfected with GSDMD-p30 (**d**) or GSDMD-FL and Caspase-1 (**e**) or GSDMD-FL and Caspase-4 (**f**), together with HDAC4-WT or its mutants. **g** Acetylation of GSDMD in HEK293T cells transfected with Flag-GSDMD and Myc-HDAC4s. **h-j** LDH release of HEK293T cells transfected with GSDMD-p30 (**h**) or GSDMD-FL and Caspase-1 (**i**) or GSDMD-FL and Caspase-4 (**j**), together with HDAC4-WT or HDAC4-3SE. ***stands for P<0.001, ** stands for P<0.01, * stands for P<0.05 and ns stands for no significant difference (unpaired t test). Data shown are mean ± SD from one representative experiments performed in triplicate.

### Protein phosphatase 1 regulates phosphorylation of HDAC4

As the phosphorylation of HDAC4 was relevant to its ability of deacetylation, we sought to find the phosphatase involved in the process. It is known that Protein phosphatase 1 (PP1) is responsible for the regulation of GSDMD phosphorylation and inhibits pyroptosis^27^. Given that HDAC4 also inhibits pyroptosis, we wondered whether PP1 interacts with HDAC4. We first confirmed that PP1α and PP1γ interacted with GSDMD (Supplementary Fig. 8a). Then we tested the interaction between PP1 and HDAC4. The results showed that HDAC4 interacted with PP1α and PP1γ, just the same as GSDMD (Fig. 6a). Immunofluorescence analysis also showed that PP1 and HDAC4 co-localized well in cells (Supplementary Fig. 8b). Structural modeling results showed that they could bind well and there were mainly hydrogen bonds and van der Waals contact between them (Supplementary Fig. 8c-e). We further explored the interaction of HDAC4, GSDMD and PP1. Immunofluorescence analysis showed that they mainly co-localized in cytoplasm (Fig. 6i). We observed that GSDMD and HDAC4 formed two hydrogen bonds, while two salt bridges and three hydrogen bonds were formed with PP1 through structural modeling (Fig. 6b-c). Then we used Phos-tag SDS-PAGE to detect the phosphorylation levels of HDAC4 with or without PP1. In the gel containing Phos-tag, the phosphorylated proteins specifically bind to the Phos-tag reagent and the migration speed of the proteins decreases as the degree of phosphorylation of the protein increases. HDAC4 co-transfection with PP1α and PP1γ showed less mobility shift (Fig. 6d), suggesting that HDAC4 is dephosphorylated by PP1α and PP1γ. THP-1 cells treated with okadaic acid (OA, a PP1 inhibitor) were subject to anti-HDAC4 immunoprecipitation and showed more mobility shift (Fig. 6e). To investigate the effect of PP1 on HDAC4-mediated pyroptosis inhibition, HEK293T cells were first co-transfected with HDAC4 and PP1α or PP1γ for 24 h, followed by transfection of GSDMD-p30 or GSDMD-FL and Caspase-1 or GSDMD-FL and Caspase-4. The results showed that PP1γ could counteract the inhibitory effect of HDAC4 on pyroptosis (Fig. 6f-h). Moreover, OA treatment could promote further inhibition of HDAC4 on pyroptosis (Supplementary Fig. 8f-h). Collectively, the above results indicate that PP1 participates in the processing of GSDMD deacetylation by dephosphorylating HDAC4.

**Fig. 6.**
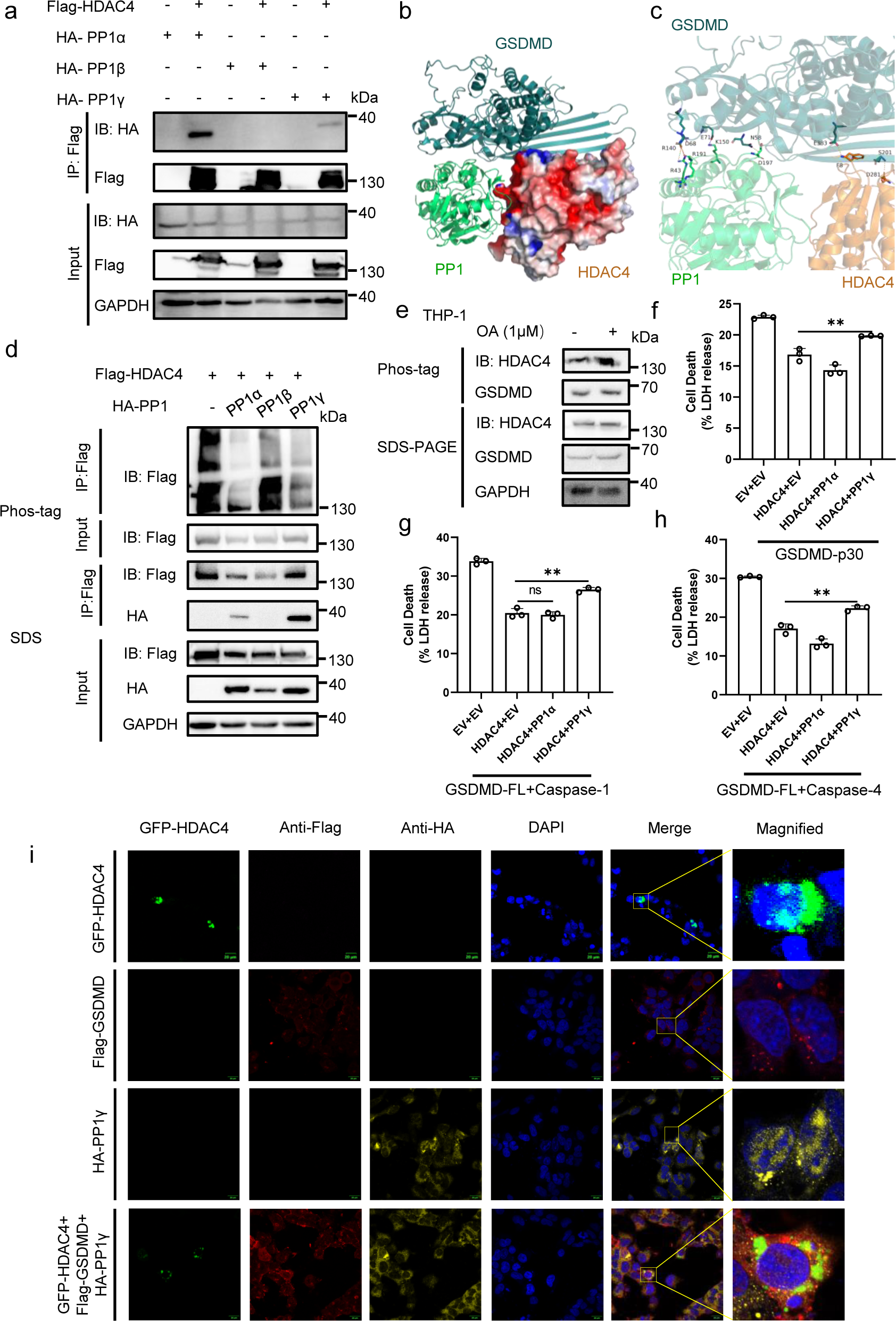
PP1 dephosphorylates HDAC4. **a** Interaction of HDAC4 and PP1. IB of total cell lysates (input) and proteins immunoprecipitated with anti-Flag resin from HEK293T cells transfected with Flag-HDAC4 and HA-PP1. **b-c** Structural model of the complex of GSDMD, HDAC4 and PP1 (PDB: 4MOV). The docking models were analyzed using HDOCK Server. **d** Analysis of phosphorylation of HDAC4 in HEK293T cells transfected with Flag-GSDMD and HA-PP1. The cells were lysed, and their proteins were separated by Phos-tag PAGE and subjected to Western blot analysis. **e** THP-1 cells were treated with or without Okadaic acid (OA) and then lysed. Proteins were separated by Phos-tag PAGE and subjected to Western blot analysis. **f-h** HEK293T cells were first transfected with HDAC4 and PP1. After 24 h, cells were then transfected with GSDMD-p30 (**f**) or GSDMD-FL and Caspase-1 (**g**) or GSDMD-FL and Caspase-4 (**h**). The supernatants were collected for LDH assay. **i** Immunofluorescence microscopy and nuclear staining (with the DNA-binding dye DAPI) of HEK293T cells transfected with expression plasmids for GFP-HDAC4, Flag-GSDMD and HA-PP1γ. Scale bars, 20 μm. ** stands for P<0.01 and ns stands for no significant difference (unpaired t test). Data shown are mean ± SD from one representative experiments performed in triplicate.

### Acetylation of GSDMD facilitates the ubiquitination of GSDMD

Protein acetylation and ubiquitination are often associated. Cross-talk between lysine acetylation and ubiquitination have been proved to be an important regulatory mechanism in regulating protein functions^15,18,19,28-30^. We first checked whether the acetylation of GSDMD-p30 affected its oligomerization. The results showed that HDAC4 transfection or TSA-treatment did not affect GSDMD-p30 oligomerization (Supplementary Fig. 9a). Furthermore, we confirmed that HDAC4 did not impact on the degradation of GSDMD (Supplementary Fig. 9b). We also detected the binding of GSDMD-WT or GSDMD-mutants and HDAC4, and the results showed that the binding was not affected (Supplementary Fig. 9c). Then we turned to investigate the relationship between acetylation and ubiquitination of GSDMD. TSA-treatment promoted the ubiquitination of GSDMD (Fig. 7a). Moreover, HDAC4 could inhibit GSDMD ubiquitination, while HDAC-H803L and HDAC4-3SA could not (Fig. 7b). The ubiquitination level of GSDMD in HEK293T cells transfected with sg-HDAC4 were less than cells transfected with NC (Fig. 7c). Then we constructed the acetylation-mimetic mutants GSDMD-K103Q and GSDMD-K248Q and examined their ubiquitination level in HEK293T cells. Compared with GSDMD-WT, GSDMD-K248Q displayed higher extent of ubiquitylation (Fig. 7d). As we have proved that GSDMD could be ubiquitinated with K27-linked polyubiquitin chains^16^, we next investigated whether the acetylation of GSDMD influences K27-linked ubiquitination of GSDMD and found that GSDMD-K248Q also had greater ubiquitination level (Fig. 7e). Thus, these results suggest that GSDMD acetylation facilitates its ubiquitination, which promotes GSDMD-mediated pyroptosis.

**Fig. 7.**
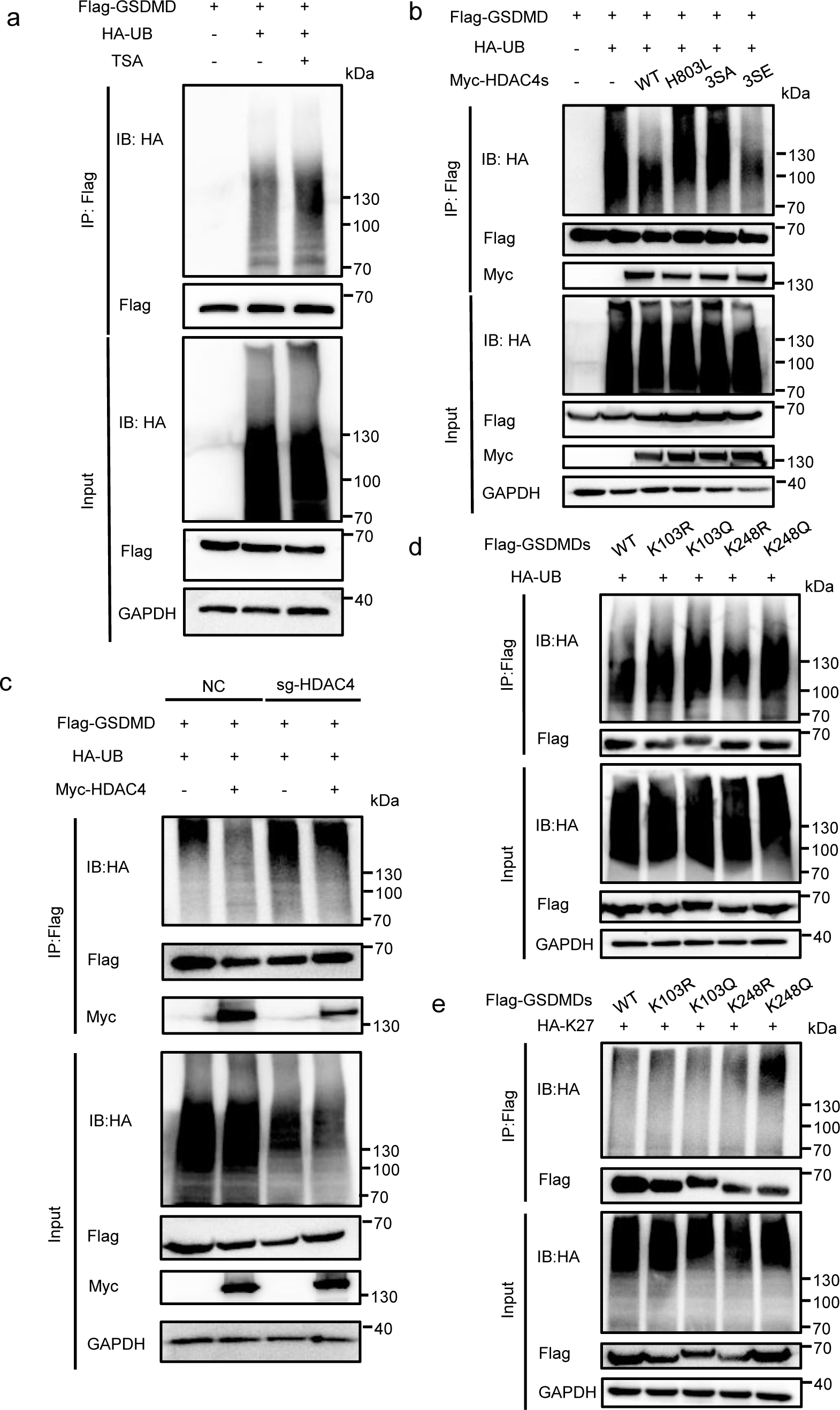
Acetylation of GSDMD promotes its ubiquitination. **a** IB of total cell lysates (input) and proteins immunoprecipitated with anti-Flag resin from HEK293T cells transfected with Flag-GSDMD and HA-UB, treat with or without TSA. **b** IB of total cell lysates (input) and proteins immunoprecipitated with anti-Flag resin from HEK293T cells transfected with Flag-GSDMD and HA-UB, together with HDAC4 or its mutants. **c** NC and sg-HDAC4 HEK293T cells were transfected with Flag-GSDMD, together with or without Myc-HDAC4. GSDMD ubiquitination was analyzed by immunoprecipitation with an anti-GSDMD antibody followed by western blotting. **d-e** IB of total cell lysates (input) and proteins immunoprecipitated with anti-Flag resin from HEK293T cells transfected with Flag-GSDMD and HA-UB (**d**) or HA-K27(**e**). All results shown are representative of at least three independent experiments.

## Discussion

GSDMD acts downstream of inflammasome activation and is responsible for forming transmembrane pores and cytokine secretion. However, the regulation of GSDMD remains to be uncovered. We have previously demonstrated that GSDMD is modified by ubiquitination, thereby promoting pyroptosis^16^. Occasionally, we also found some acetyltransferases and the deacetylases may also be involved in the ubiquitination of GSDMD. Thus, we reveal a previously unknown post-translational modification of GSDMD. GSDMD can be acetylated in the resting state and the acetylation level of GSDMD increases after NLRP3 inflammasome activation. Subsequently, acetylation of GSDMD promotes pyroptotic cell death.

Acetylation was initially described as an epigenetic mechanism modulating nucleosomal DNA accessibility and transcription. After then, acetylation was shown to be an essential key to regulate physiological processes such as cell cycle, DNA damage repair, and cellular signalling^31-34^. Acetylation of proteins has effects on protein-protein interactions and protein degradation. Both proteasome-dependent and proteasome-independent protein degradation can be affected by acetylation. PEPCK1 acetylation results in PEPCK1 ubiquitination and degradation, while acetyltransferases CBP, p300 and TIP60 and deacetylases HDAC6 and SIRT1 are important regulators of autophagy^35,36^. In this study, we found that GSDMD was acetylated at Lys 248. We also identified HDAC4 as the deacetylase of GSDMD. HDAC4 directly interacted with GSDMD in multiple sites and deacetylates GSDMD. Intriguingly, we found that the H803 site of HDAC4, responsible for its deacetylation activity, had no effect on inhibiting pyroptosis. However, the S246, S467 and S632 residues of HDAC4 played important roles in the process of deacetylation and suppressing pyroptosis. We speculated that the three sites affected the export of HDAC4 to cytoplasm, where it bound to GSDMD and functioned. HDAC4 is reported to be phosphorylated by the signaling kinase TBK1/IKKε, thus preventing IRF3 phosphorylation^26^. But the phosphate mediating the dephosphorylation of HDAC4 is not clear for now. PP1 is found to dephosphorylate GSDMD^27^. Interestingly, we found that PP1 also bound with HDAC4, together with GSDMD, and dephosphorylated HDAC4. Thus, PP1 itself could inhibit pyroptosis by dephosphorylating GSDMD, while it also affects the deacetylase activity of HDAC4 on GSDMD through inhibiting HDAC4 phosphorylation, thereby inhibiting the translocation of HDAC4 to cytoplasm.

Proteins are modified by various PTMs, which can reciprocally influence each other. The lysine residues can be diversely modified, including acetylation, methylation and ubiquitination, which could lead to competitive PTMs crosstalk whereby different PTMs compete for the same residue. For example, acetylation of p53 competes with ubiquitination mediated by MDM2 on the same lysine residues, thereby inhibiting its degradation through ubiquitin-proteasome pathway^37^. However, PTMs do not always behave competitively. Acetylation at K420 and K435 sites enhances p62 binding to ubiquitin and K435 acetylation also directly increases the UBA-ubiquitin affinity^19^. Our results indicated that the acetylation of GSDMD at K248 site promotes GSDMD K27-linked ubiquitination, which provides a new example for crosstalk between acetylation and ubiquitination.

In summary, we have revealed an unknown posttranslational modification of GSDMD and a novel interplay among GSDMD, HDAC4 and PP1 (Fig. 8). The PTMs regulations of GSDMD and HDAC4, including the ubiquitination and acetylation of GSDMD and the phosphorylation of HDAC4, modulate the pyroptosis. These mechanistic studies could provide insights into the mechanism underlying pyroptosis and offer an opportunity to intervene inflammatory-related diseases.

**Fig. 8.**
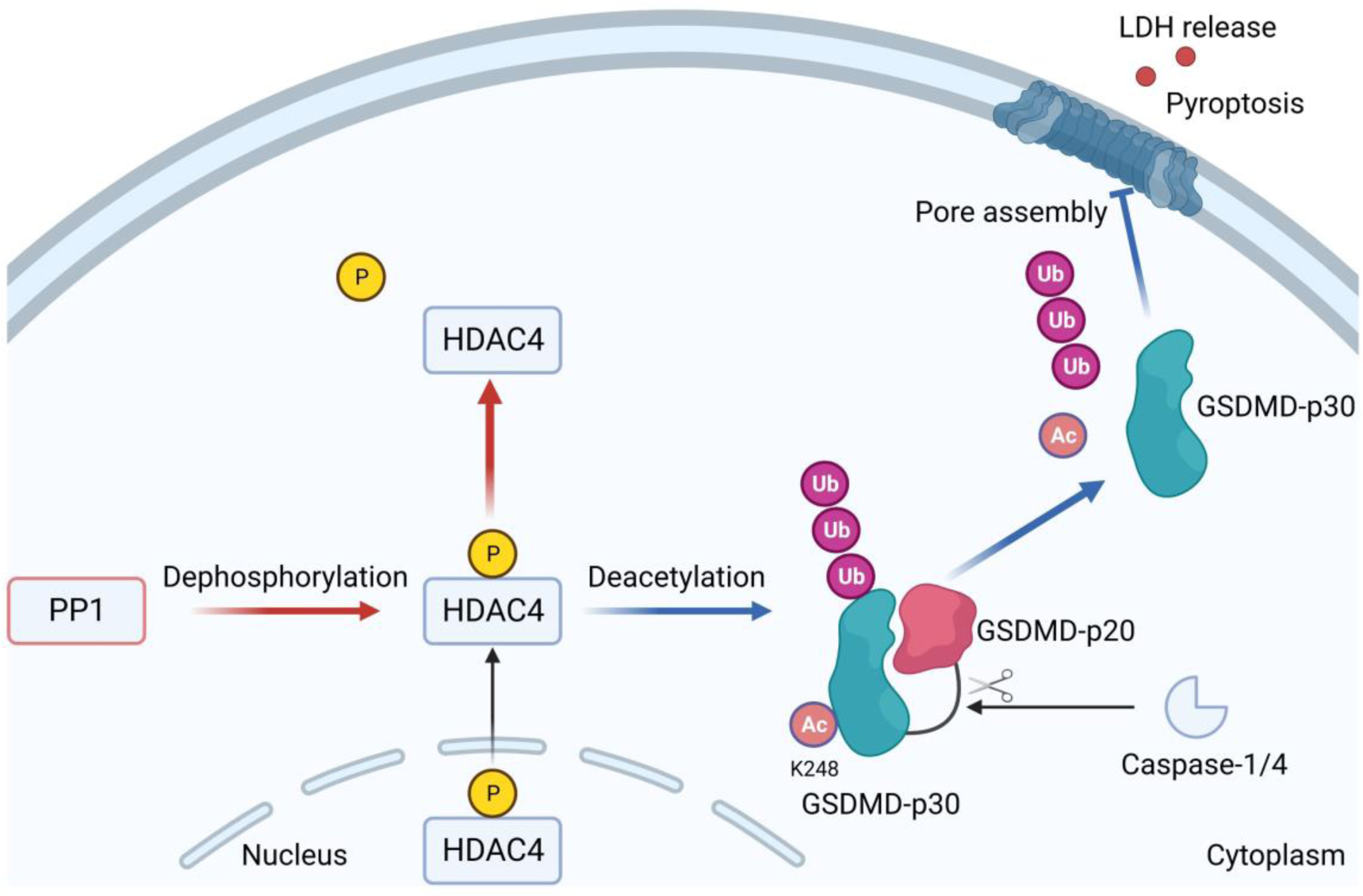
Diagram of HDAC4 regulates GSDMD-mediated pyroptosis.

## Methods

### Reagents and antibodies

Anti-HA (3724), anti-GSDMD (39754) and anti-Acetylated Lysine (9441) antibodies were purchased from Cell Signaling Technology. Anti-GAPDH antibody, HRP-labeled goat anti-mouse IgG and goat anti-rabbit IgG were from Hangzhou Fudebio. Anti-Flag M2 (F1804), anti-Flag M2 Magnetic Beads (10004D), anti-Myc (C3956) antibodies and LPS (O111:B4, L2630) were purchased from Sigma-Aldrich. Anti-FLAG antibody (rabbit source), Goat pAb to MS IgG (Chromeo 546, ab60316), Dnk pAb to Rb IgG (Alexa Fluor 647, ab150075) fluorescent secondary antibodies were obtained from Abcam. Nigericin, Pam3CSK4 (TLRL-PMS), poly(dA:dT) (TLRL-PATN) and flagellin (TLRL-EPSTFLA) were purchased from InvivoGen. ELISA kits for analysis of human IL-1β, mouse IL-1β, mouse IL-6 and mouse TNF-α were purchased from MultiSciences. CytoTox 96 LDH-release assay kit (G1780) was from Promega. Efficient eukaryotic transfection reagent VigoFect was purchased from Vigorous Biotechnology (Beijing), Lipofectamine 2000 transfection reagent was obtained from Invitrogen while Lipo8000^TM^ transfection reagent was obtained from Beyotime Biotechnology. Trichostain A (HY-15144), Nicotinamide (HY-B0150), LMK-235(HY-18998) and okadaic acid (HY-N6785) were purchased from MCE.

### Plasmid construction and transfection

Flag-GSDMD, Flag-GSDMD-p30, HA-GSDMD-p30, HA-Caspase-1, Myc-Caspase-4 and HA-tagged ubiquitin was constructed previously. Myc-HDAC4 was made by cloning human HDAC4 protein ORF (GeneBank: NM_001378415.1) into a pCMV-Myc vector using NotI and SalI restriction sites. HDAC4 mutants were constructed based on the eukaryotic expression plasmids Myc-HDAC4. The primers used in this study are shown in Supplementary Table 1. According to the manufacturer’s instructions, indicated plasmids were transfected using VigoFect, Lipofectamine 2000 or Lipo8000 separately.

### Cell culture and stimulation

All cells were cultured at 37 °C in 95% air and 5% CO _2_. HEK293T cells, IPEC-J2 cells, RAW264.7 cells and NIH-3T3 cells were cultured in Dulbecco’s modified Eagle’s medium (DMEM) with 10% FBS (Excell) and 1% penicillin-streptomycin (Hyclone). Human monocyte cell line THP-1 cells were cultured in RPMI 1640 medium with added 10% FBS (Gibco) and 1% penicillin-streptomycin (Hyclone). To induce inflammasome activation, 5 × 10^5^ THP-1 cells were plated in a 24-well plate overnight with 1 μM PMA, and then the medium was changed next morning. In order to activate the canonical NLRP3 inflammasome, cells were primed for 4 h with 500 ng/mL LPS and then stimulated for 1 h using 10 μM Nigericin. For activation of the non-canonical inflammasome, cells were primed for 4 h with 250 ng/mL Pam3CSK4, after which the medium was replaced and cells were transfected with 2 μg/mL LPS for 6 h. For activation of AIM2 inflammasome and NLRC4 inflammasome, cells were primed for 4 h with 500 ng/mL LPS, followed by stimulating for 6 h with 2 μg/mL poly(dA:dT) or flagellin.

### Immunoblotting

Cells were harvested and lysed in RIPA lysis buffer (Beyotime Biotechnology) added with 1% PMSF (Beyotime Biotechnology). Proteins were separated on the 10% SDS-PAGE gel (Hangzhou Fudebio) and transferred onto the PVDF membranes (Bio-rad). Membranes were blocked in the blocking buffer (Beyotime Biotechnology), followed by staining with primary antibodies and positioning with secondary antibodies. Chemiluminescent signals were captured by an ECL chemiluminescence imaging analysis system (Clinx Science Instruments).

### Co-immunoprecipitation

For GSDMD acetylation analysis, cells were lysed in IP lysis buffer (Beyotime Biotechnology) added with PMSF. Then the cell lysate was mixed with antibodies at 4 °C overnight followed by t he addition of protein G beads (10003D, Invitrogen). Otherwise, cells were lysed in IP lysis buffer added with PMSF. The supernatants were then incubated with anti-Flag binding beads (Sigma, M8823) at 4 °C. The immunocomplexes were washed and then subjected to immunoblotting analysis.

### Confocal immunofluorescence assay

HEK293T cells were seeded on coverslips in 24-well plates. After being transfected for 24 h, cells were fixed with Immunol Staining Fix Solution (Beyotime). Cells were incubated with primary antibodies overnight at 4 °C after being permeabilized and blocked. Alex Fluor 546/647-conjugated secondary antibody was incubated for 1 h. Nuclei were stained with DAPI. Confocal micrographs were imaged using a laser confocal microscope (Olympus).

### Cytotoxicity assay

The supernatants of cells were collected and then applied to cytotoxicity test using CytoTox 96^®^ Reagent (Promega) according to the manufacturer’s manual. OD values were collected at 492 nm on an enzyme marker (Thermo Scientific).

### ELISA

Supernatants from stimulated cells, mice serum and peritoneal lavage fluid were applied to the detection of IL-1β/TNF-α/IL-6 according to the manufacturer’s instructions. Each trial group was conducted independently for three times.

### Phos-tag SDS-PAGE

Supplements of 50 μM phos-tag and 10 mM MnCl_2_ were added during preparation of SDS-PAGE gel. Others were same as above protocol of immunoblotting.

### Construction of HDAC4 knockout cells

HDAC4 knockout HEK293T cells were generated using the CRISPR/Cas9 technique. Vectors expressing gRNA targeting human HDAC4 (5’-3’ TGTTCTCCAGATGGACTTTC) were transfected into HEK293T cells. For HDAC4 knockout THP-1 cells, gRNA plasmid was co-transfected with the lentiviral packaging vectors pMD2G and PSPAX2, then introduced into HEK293T cells to produce lentivirus. After 48 h, the viral supernatants were collected and added to THP-1 cells in 6-well plates with 1 mL medium containing 8 μg/mL polybrene. The infected cells were spun at 300 ×g for 1 h, and then 1 mL fresh media was added. After infection for 2 days, stably transfected cells were selected with GFP by flow-cytometric analysis. Single-cell sorting of transfected cells was performed using flow cytometry (MOFLO XDP).

### RNA interference

SiRNAs (Genepharma) specific for mouse HDAC4 were transfected into NIH-3T3 cells using the GP-transfect-Mate according to the manufacturer’s instructions. Then the si-HDAC4#1 (5’-3’ CACCAUCCUUACCCAACAUTT) were injected into mice (50 μg /mouse) in the presence of in vivo RNA transfection reagent (Entranster-in vivo; Engreen). siRNA sequences are shown in Supplementary Table 2.

### h-GSDMD reconstruction lentivirus

The lentivirus was produced by transfection of the pLVX-IRES-Puro-h-GSDMD-WT/ h-GSDMD-K248R or control vector into HEK293T cells using Lipo8000 with pMD2G and PSPAX2. Supernatants were collected after 72 h incubation for further lentiviral concentration.

### LC-MS/MS analysis

Flag-tagged GSDMD immunoprecipitants prepared from whole-cell lysates or gel-filtrated fractions were resolved on SDS-PAGE gels, and protein bands were excised. The samples were digested with trypsin, and then subject to LC-MS/MS analysis. Swissprot_Human mass spectra were used as the standard reference. Trypsin/P was used for cleavage. MS data were captured and analyzed by Shanghai Bioprofile.

### Ethics statement

Animal experiments were conducted following guidelines for experimental animals’ welfare and ethics. All animal studies were reviewed and approved by Laboratory Animal Center of Zhejiang University.

### Statistical analysis

All experiments were performed independently at least three times. Data were presented as the mean ± standard deviation (SD), analyzed and used for statistical graphing by GraphPad Prism 8, the significance of differences was determined by Student’s *t*-test. The significance of differences ranked as: ***stands for P<0.001, ** stands for P<0.01, * stands for P<0.05 and ns stands for non-significant difference.

## Data availability

The authors declare that all data supporting the findings of this article are available within the paper and the supplementary information files or are available from the authors upon request.

## Acknowlegements

This work was financially supported by the National Key Research and Development Program of China (2022YFD1800300), the National Natural Science Foundation of China (32072817), the Zhejiang Provincial Key R&D Program of China (2021C02049), the Joint Fund of the National Natural Science Foundation of China and Guangdong province (U22A20521), the “Pioneer” and “Leading Goose” R&D Program of Zhejiang Province (2022C02031), High-level Talents Special Support Plan of Zhejiang Province (2021R52041), the Agricultural Major Technology Synergy Extension Project of Zhejiang Province (2021XTTGXM02-05), the Scientific Research Fund of Zhejiang University (XY2022004) and the Fundamental Research Funds for the Central Universities (2022-KYY-517101-0005).

We thank Dr. Weiren Dong and Ying Shan in the Shared Experimental Platform for Core Instruments, College of Animal Sciences, Zhejiang University for assistance with the analysis of laser confocal microscopy imaging. We thank Mr. Baochun Su and Zhipeng Ma from Jinrong Peng Lab for kindly providing NIH-3T3 cells.

## Author contributions

F.S.S., W.L.X. and Y.Y conceived and designed the study. W.L.X, X.Y.L, D.Y.L, X.Y.F, N.C, Q.L, Y.H.S, S.H.H and L.D performed the experiments. W.L.X and F.S.S wrote the paper. All authors have contributed to, reviewed, and approved the manuscript.

