## Supplementary Information for "HDAC4-mediated deacetylation of GSDMD prevents pyroptosis by inhibiting GSDMD ubiquitination"

Weilv *et al*.

### Supplementary Information

### Inventory of Supporting Information

### 1. Supplementary Figures and figure legends

**Supplementary Fig. 1** PI staining for Fig. 1g-i.

**Supplementary Fig. 2** TSA-treatment promotes pyroptosis.

**Supplementary Fig. 3** TSA and LMK-235 have no effect on cytokine secretion.

**Supplementary Fig. 4** Candidate acetylation sites of GSDMD.

**Supplementary Fig. 5** HDAC4 suppresses pyroptotic cell death.

**Supplementary Fig. 6** PI staining for Fig. 3f.

**Supplementary Fig. 7** LMK-235-treatment promotes pyroptosis.

**Supplementary Fig. 8** PP1 mediated dephosphorylation of HDAC4.

**Supplementary Fig. 9** HDAC4 does not affect GSDMD oligomerization or degradation.

### 2. Supplementary Tables

**Supplementary Table 1.** Primers used in this study for the construction of plasmids.

**Supplementary Table 2.** siRNA sequences for the mouse HDAC4 oligonucleotide.

### 1. Supplementary Figures and figure legends

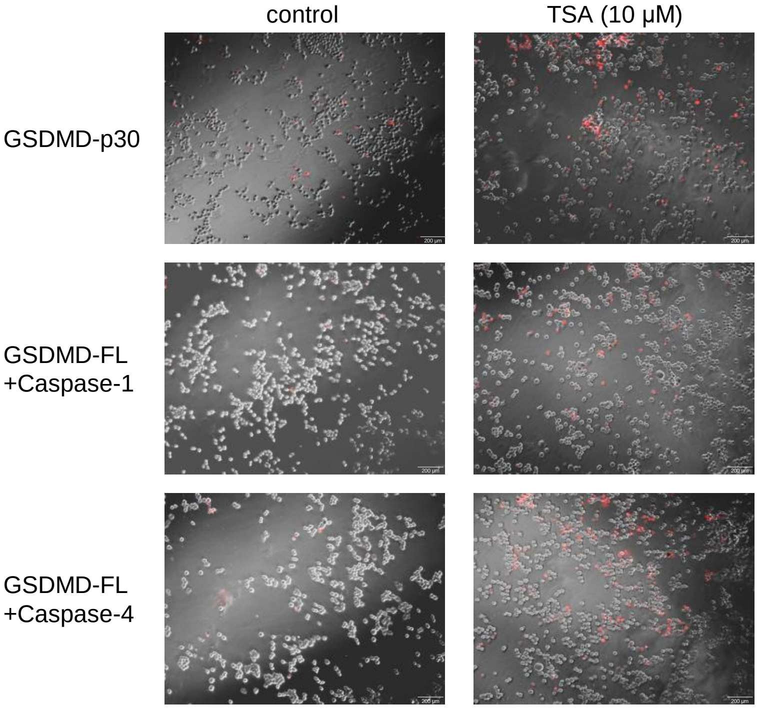


**Supplementary Fig. 1 PI staining for Fig. 1g-i.** PI staining of HEK293T cells after 24 h transfection with GSDMD-p30 or GSDMD-FL and Caspase-1 or GSDMD-FL and Caspase-4, treated with or without 10 μM TSA. Scale bars, 200 μm.

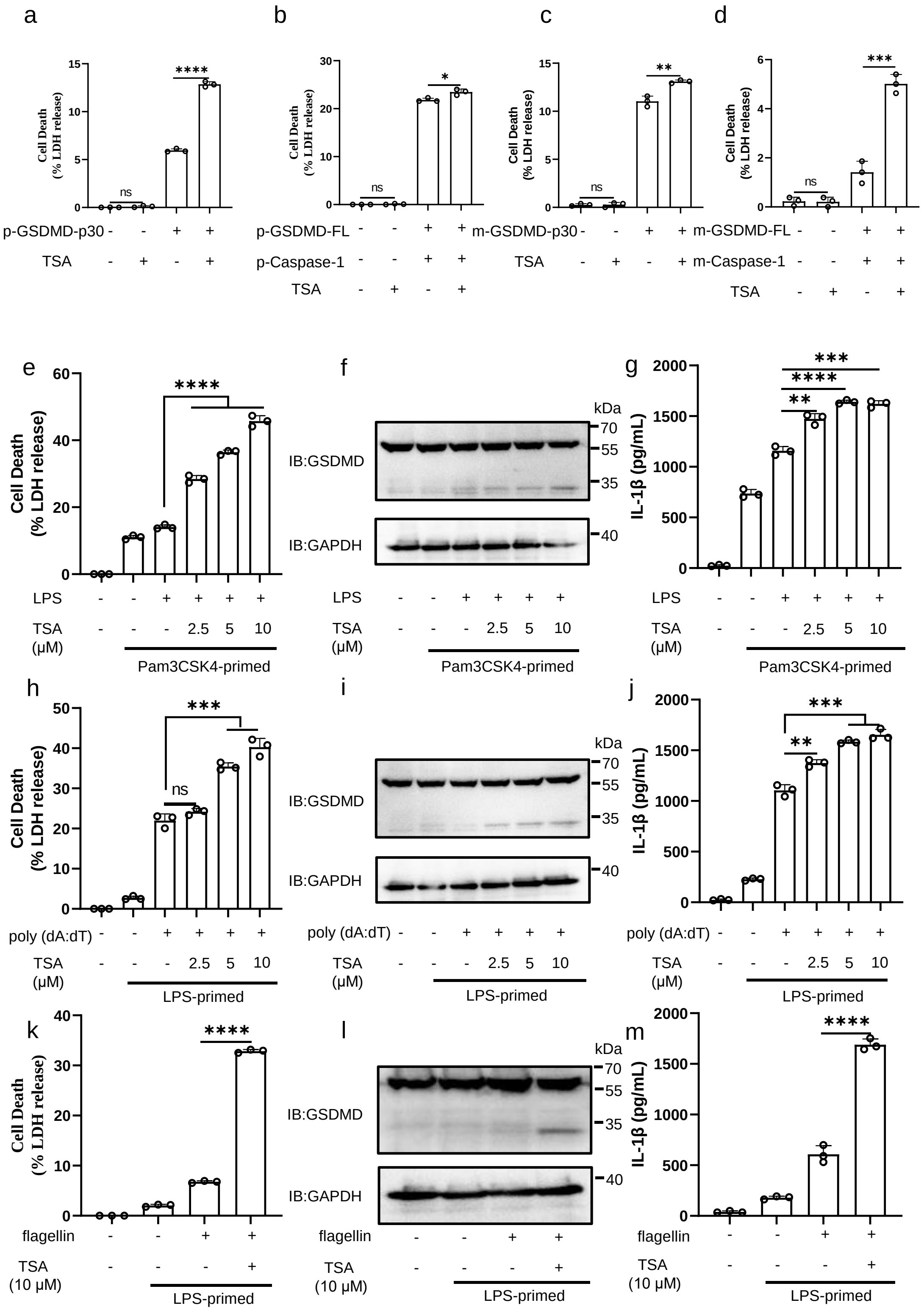


**Supplementary Fig. 2 TSA-treatment promotes pyroptosis. a-d** HEK293T cells were transfected with pig (**a**, **b**) or mouse (**c**, **d**) GSDMD-p30 or GSDMD-FL and Caspase-1, treat with or without 10 μM TSA. The supernatants were collected for

LDH assay. **e-g** PMA-differentiated THP-1 cells were primed with 250 ng/mL Pam3CSK4 and TSA for 4 h, and then transfected with 2 μg/mL LPS for 6 h. The supernatants were collected and analyzed by LDH release assay (**e**) and ELISA for IL-1β (**g**). Cell lysates were analyzed by immunoblotting (**f**). **h-j** PMA-differentiated THP-1 cells were primed with 500 ng/mL LPS and TSA for 4 h, and then transfected with 2 μg/mL poly(dA:dT) for 6 h. The supernatants were collected and analyzed by LDH release assay (**h**) and ELISA for IL-1β (**j**). Cell lysates were analyzed by immunoblotting (**i**). **k-m** PMA-differentiated THP-1 cells were primed with 500 ng/mL LPS and TSA for 4 h, and then transfected with 2 μg/mL flagellin for 6 h. The supernatants were collected and analyzed by LDH release assay (**k**) and ELISA for IL-1β (**m**). Cell lysates were analyzed by immunoblotting (**l**). $^{****}$stands for P<0.0001, $^{***}$stands for P<0.001, $^{**}$ stands for P<0.01, $^{*}$stands for P<0.05 and ns stands for no significant difference (unpaired t test). Data shown are mean ± SD from one representative experiments performed in triplicate.

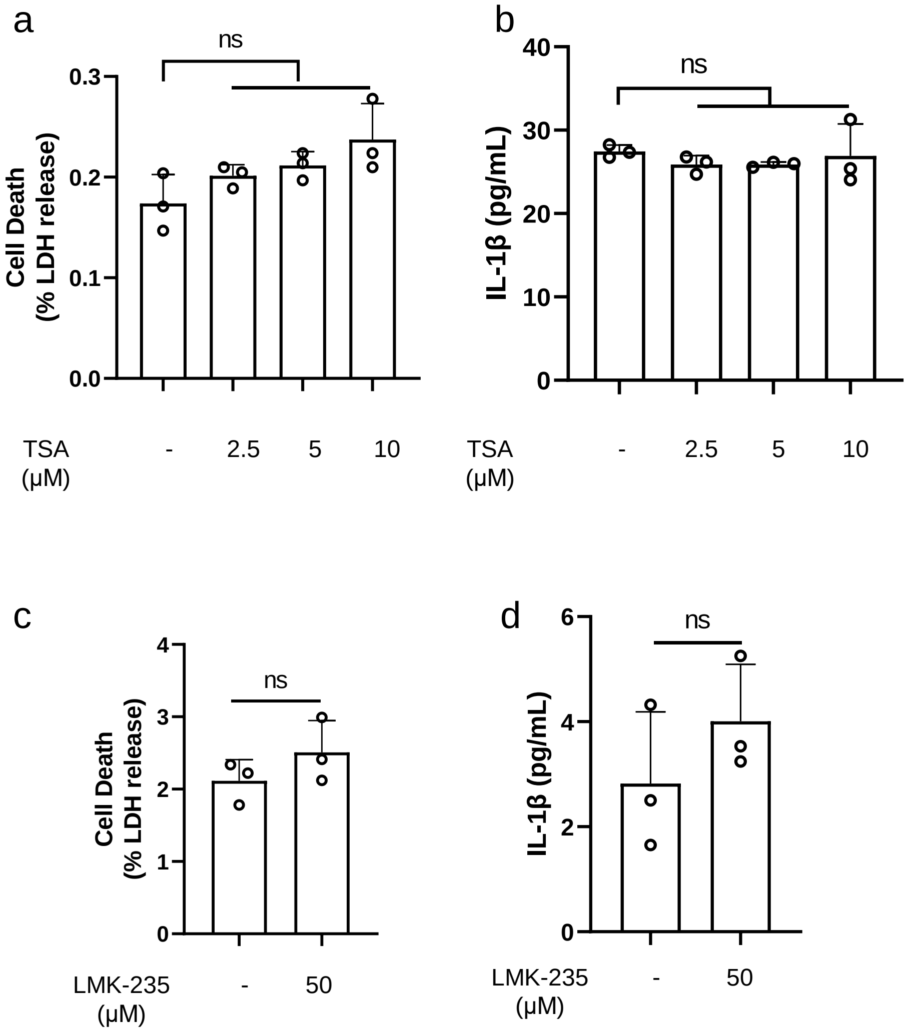


**Supplementary Fig. 3 TSA and LMK-235 have no effect on cytokine secretion.** **a-b** THP-1 cells were treated with TSA for 6 h, and supernatants were collected and analyzed by LDH release assay (**a**) and ELISA for IL-1β (**b**). **c-d** THP-1 cells were treated with LMK-235 for 6 h, and supernatants were collected and analyzed by LDH release assay (**c**) and ELISA for IL-1β (**d**). Ns stands for no significant difference (unpaired t test). Data shown are mean ± SD from one representative experiments performed in triplicate.

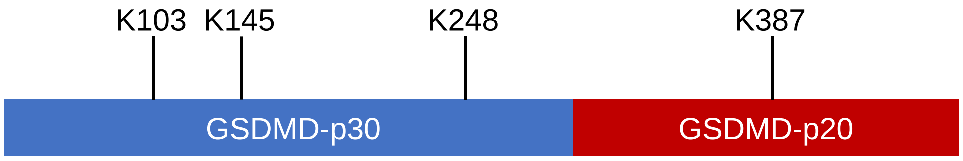


MGSAFERVVRRVVQELDHGGEFIPVTSLQSSTGFQPYCLVVRKPSSSWFW
KPRYKCVNLSIKDILEPDAAEPDVQRGRSFHFYDAMDGQIQGSVELAAPGQ
AKIAGGAAVSDSSSTSMNVYSLSVDPNTWQTLLHERHLRQPEHKVLQQLRS
RGDNVYVVTEVLQTQKEVEVTRTHKREGSGRFSLPGATCLQGEGQGHLSQ
KKTVTIPSGSTLAFRVAQLVIDSDLDVLLFPDKKQRTFQPPATGHKRSTSEGA
WPQLPSGLSMMRCLHNFLTDGVPAEGAFTEDFQGLRAEVETISKELELLDR
ELCQLLLEGLEGVLRDQLALRALEEALEQGQSLGPVEPLDGPAGAVLECLVL
SSGMLVPELAIPVVYLLGALTMLSETQHKLLAEALESQTLLGPLELVGSLLEQ
SAPWQERSTMSLPPGLLGNSWGEGAPAWVLLDECGLELGEDTPHVCWEP
QAQGRMCALYASLALLSGLSQEPH

| | 103 | 145 | 248 | 387 |
|---|---|---|---|---|
| Human | 99 GQAKIAGG | 142 PEHKVLQ | 245 GHKRST | 384 TQHKLLA |
| Mouse | 100 GEGKISGG | 143 PENKILQ | 246 DRKAVG | 387 TQQQLLA |
| Rattus | 100 GQGKISGG | 143 PENKILQ | 246 GHKGAV | 387 TQQQLLA |
| Pig | 99 GQGKFSGK | 142 PEHKVLQ | 245 GHSASH | 388 TQHVLLA |
| Cat | 99 GQGKFVSG | 142 PEHKILQ | 245 GHKPTS | 387 TQRVLLA |
| Macaca | ………………… | 25 PEHKILQ | 128 GHKPSR | 267 TQHKLLA |

**Supplementary Fig. 4 Candidate acetylation sites of GSDMD.** Scheme of human GSDMD protein marked with four candidate acetylation sites, including K103, K145, K248 and K387. Sequence alignment of the K103, K145, K248 and K387 sites within GSDMD orthologues of different species (below).

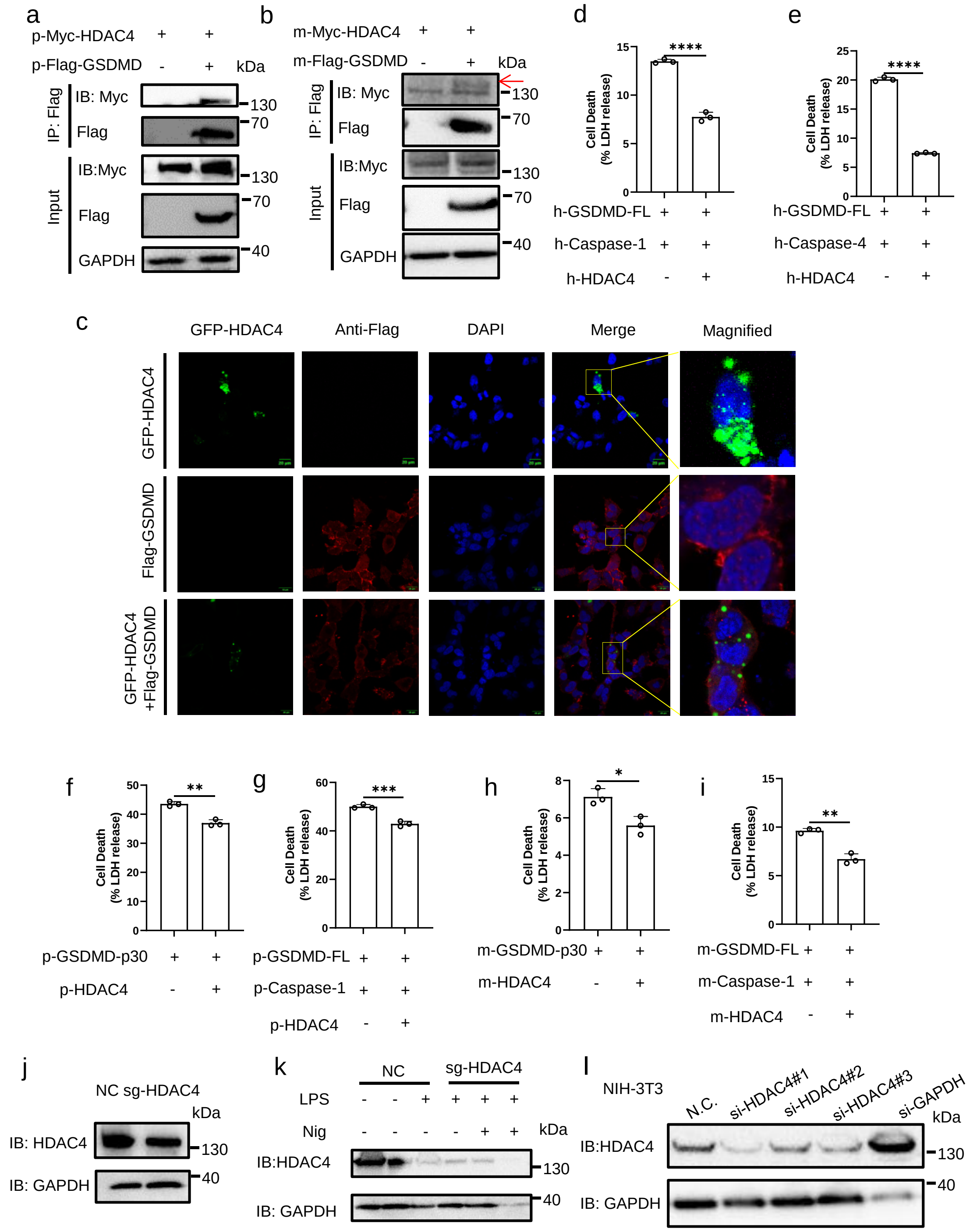


**Supplementary Fig. 5 HDAC4 suppresses pyroptotic cell death. a-b** IB of total cell lysates (input) and proteins immunoprecipitated with anti-Flag resin from HEK293T cells transfected with Myc-HDAC4 and Flag-GSDMD from pig (**a**) or mouse (**b**). **c** Immunofluorescence microscopy and nuclear staining (with the DNA-binding dye DAPI) of HEK293T cells transfected with expression plasmids for

GFP-HDAC4 and Flag-GSDMD. Scale bars, 20 μm. **d-e** LDH release of HEK293T cells transfected with GSDMD-FL and Caspase-1 or GSDMD-FL and Caspase-4, together with or without HDAC4. **f-i** LDH release of HEK293T cells transfected with GSDMD-p30 and GSDMD-FL and Caspase-1 from pig (**f-g**) or mouse (**h-i**), together with or without HDAC4. **j** IB of NC or sg-HDAC4 HKE293T cells. **k** IB of NC or sg-HDAC4 THP-1 cells treated with LPS and Nig. **l** NIH-3T3 cells were transfected with non-targeting control siRNA (si-NC) or siRNA targeting HDAC4 (si-HDAC4#1, #2, #3), or siRNA targeting GAPDH for 24 h. Cell lysates were analyzed by immunoblotting. $^{****}$stands for P<0.0001, $^{***}$stands for P<0.001, $^{**}$ stands for P<0.01, $^{*}$ stands for P<0.05 and ns stands for no significant difference (unpaired t test). Data shown are mean ± SD from one representative experiments performed in triplicate.

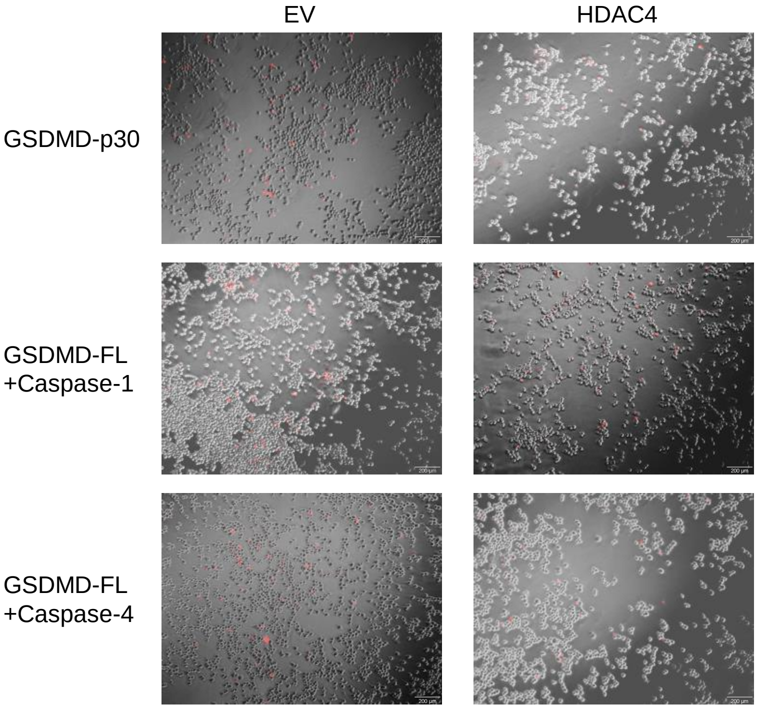


**Supplementary Fig. 6 PI staining for Fig. 3f.** PI staining of HEK293T cells after 24 h transfection with indicated plasmids. Scale bars, 200 μm.

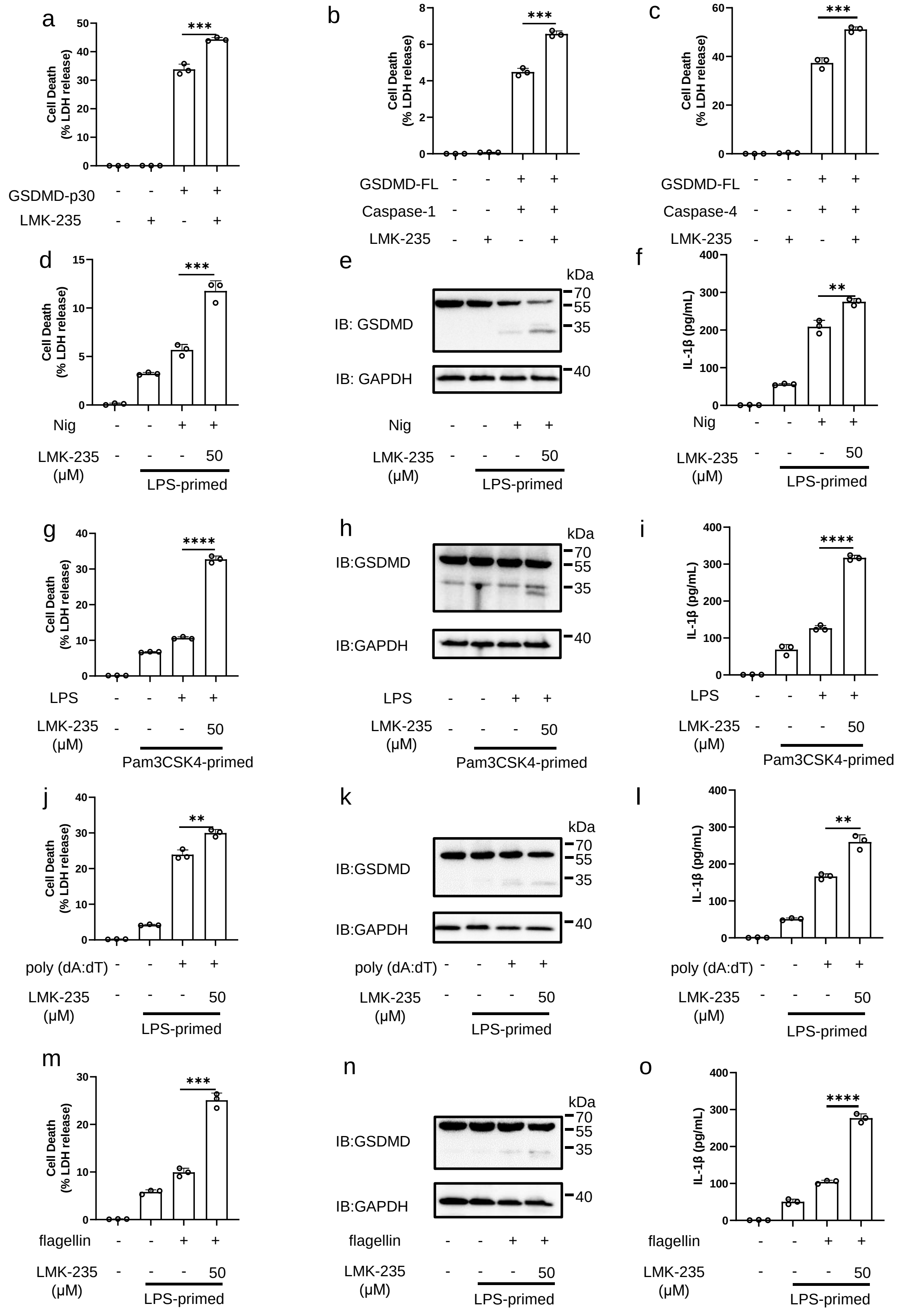


**Supplementary Fig. 7 LMK-235-treatment promotes pyroptosis. a-c** LDH release of HEK293T cells transfected with GSDMD-FL and Caspase-1 or GSDMD-FL and

Caspase-4, treated with or without LMK-235 (50 μM). **d-f** THP-1 cells were incubated with 500 ng/mL LPS and LMK-235 for 4 h and then another 1 h for 10 μM Nigericin. The supernatants were collected and analyzed by LDH release assay (**d**) and ELISA for IL-1β (**f**). Cell lysates were analyzed by immunoblotting (**e**). **g-i** PMA-differentiated THP-1 cells were primed with 250 ng/mL Pam3CSK4 and LMK-235 for 4 h, and then transfected with 2 μg/mL LPS for 6 h. The supernatants were collected and analyzed by LDH release assay (**g**) and ELISA for IL-1β (**i**). Cell lysates were analyzed by immunoblotting (**h**). **j-l** PMA-differentiated THP-1 cells were primed with 500 ng/mL LPS and LMK-235 for 4 h, and then transfected with 2 μg/mL poly(dA:dT) for 6 h. The supernatants were collected and analyzed by LDH release assay (**j**) and ELISA for IL-1β (**l**). Cell lysates were analyzed by immunoblotting (**k**). **m-o** PMA-differentiated THP-1 cells were primed with 500 ng/mL LPS and LMK-235 for 4 h, and then transfected with 2 μg/mL flagellin for 6 h. The supernatants were collected and analyzed by LDH release assay (**m**) and ELISA for IL-1β (**o**). Cell lysates were analyzed by immunoblotting (**n**). $^{****}$stands for P<0.0001, $^{***}$stands for P<0.001, $^{**}$ stands for P<0.01 (unpaired t test). Data shown are mean ± SD from one representative experiments performed in triplicate.

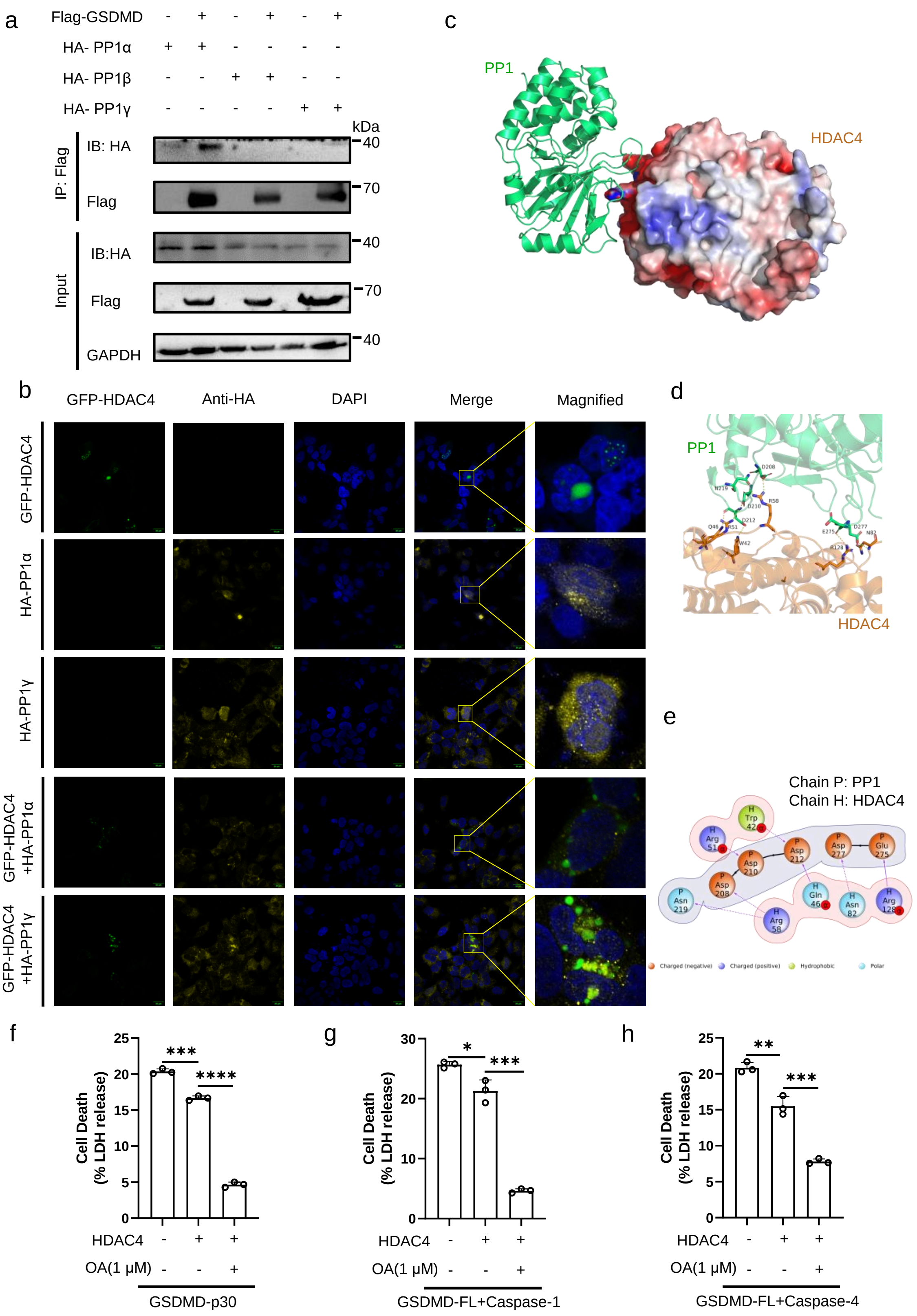


**Supplementary Fig. 8 PP1 mediated dephosphorylation of HDAC4. a** IB of total cell lysates (input) and proteins immunoprecipitated with anti-Flag resin from HEK293T cells transfected with Flag-HDAC4 and HA-PP1 subunits. **b**

Immunofluorescence microscopy and nuclear staining (with the DNA-binding dye DAPI) of HEK293T cells transfected with expression plasmids for GFP-HDAC4 and HA-PP1 subunits. Scale bars, 20 μm. **c-e** Structural model of the interaction of HDAC4 (PDB: 2VQW) and PP1 (PDB:4MOV). Hydrogen bonds are represented by yellow dotted lines in 3D display (**d**). Hydrogen bonds are represented by straight arrows or dashed lines in interaction 2D map (**e**). The docking models were analyzed using HDOCK Server. **f-h** LDH release of HEK293T cells co-transfected with GSDMD-p30 (**f**) or GSDMD-FL and Caspase-1 (**g**) or GSDMD-FL and Caspase-4 (**h**) and HDAC4, treated with or without OA (1μM). $^{****}$stands for P<0.0001, $^{***}$stands for P<0.001, $^{**}$ stands for P<0.01 and $^{*}$ stands for P<0.05 (unpaired t test). Data shown are mean ± SD from one representative experiments performed in triplicate.

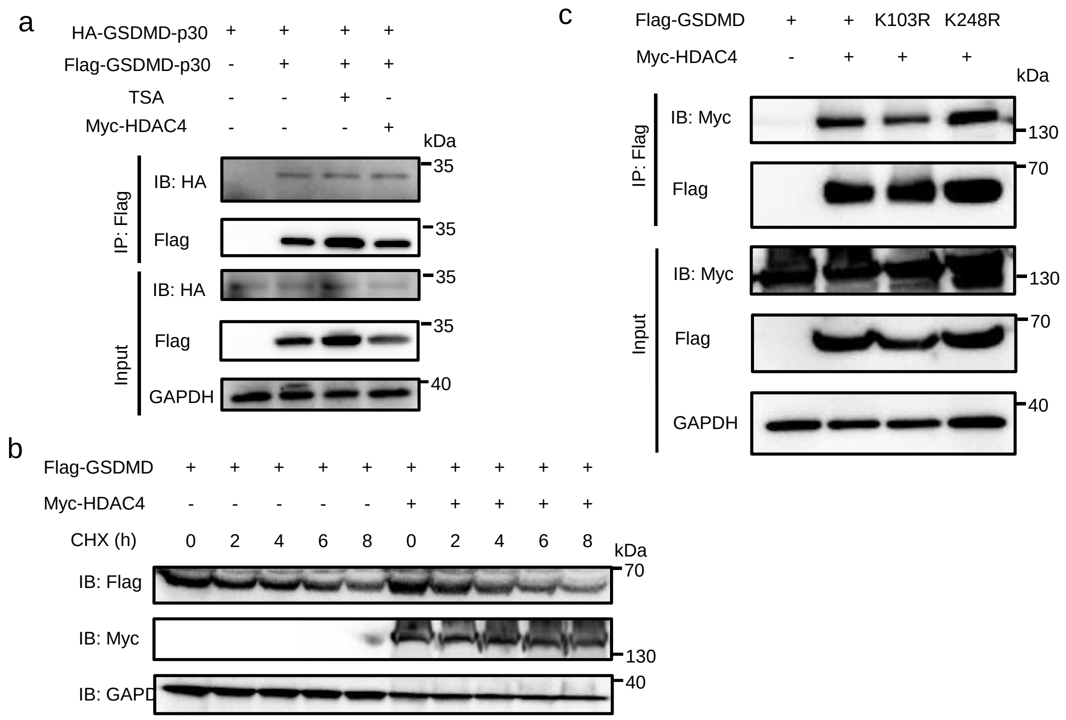


**Supplementary Fig. 9 HDAC4 does not affect GSDMD oligomerization or degradation. a** IB of total cell lysates (input) and proteins immunoprecipitated with anti-Flag resin from HEK293T cells transfected with Flag-GSDMD-p30 and HA-GSDMD-p30, together with or without Myc-HDAC4 or TSA. **b** HEK293T cells were transfected with Flag-GSDMD together with or without Myc-HDAC4, then treated with cycloheximide (CHX) (25 μg/mL) for the indicated time points. Cell lysates were analyzed by immunoblotting. **c** IB of total cell lysates (input) and proteins immunoprecipitated with anti-Flag resin from HEK293T cells transfected with Myc-HDAC4 and Flag-GSDMD or its mutants. All results shown are representative of at least three independent experiments.

### 2. Supplementary Tables

**Supplementary Table 1. Primers used in this study for the construction of plasmids.**

| **Primers** | **Sequences (5'-3')** |
|---|---|
| Myc-HDAC4-F | GGAGGCCCGAATTCGGTCGACTATGAGCTCCCAAAGCCATCC |
| Myc-HDAC4-R | CATGTCTGGATCCCCGCGGCCGCCTACAGGGGCGGCTCCTC |
| Flag-HDAC4-F | AAGGATGACGATGACAAGCTTATGAGCTCCCAAAGCCATCC |
| Flag-HDAC4-R | ATCAGATCTATCGATGAATTCCTACAGGGGCGGCTCCTC |
| GFP-HDAC4-F | TCGAGCTCAAGCTTCGAATTCAATGAGCTCCCAAAGCCATCC |
| GFP-HDAC4-R | CGGGCCCGCGGTACCGTCGACCTACAGGGGCGGCTCCTC |
| HDAC4-ΔNLS-F | GGAGGCCCGAATTCGGTCGACTATGCTGGCCATGAAGCACC |
| HDAC4-ΔNLS-R | CATGTCTGGATCCCCGCGGCCGCCTACAGGGGCGGCTCCTC |
| HDAC4-ΔN-F | GGAGGCCCGAATTCGGTCGACTGGCCTCGTGTATGACACGC |

| | |
|---|---|
| HDAC4-ΔN-R | CATGTCTGGATCCCCGCGGCCGCCTACAGGGGCGGC TCCTC |
| HDAC4-ΔC-F | GGAGGCCCGAATTCGGTCGACTATGAGCTCCCAAAG CCATCC |
| HDAC4-ΔC-R | CATGTCTGGATCCCCGCGGCCGCCTATGTCGTGAACC TCGGCTTG |
| HDAC4-ΔNES-F | GGAGGCCCGAATTCGGTCGACTATGAGCTCCCAAAG CCATCC |
| HDAC4-ΔNES-R | CATGTCTGGATCCCCGCGGCCGCCTATTCGTTCTCGC AAGTCTGAGC |
| HDAC4-H803L-F | GGAGGCCCGAATTCGGTCGACTATGAGCTCCCAAAG CCATCC |
| HDAC4-H803L-R | TCTCCTCCGCAAGGTGTCCAGGGGGGCGGACC |
| HDAC4-S246A-F | AACAGCTGCTGAACCGAATCTGAAATTACGGTCC |
| HDAC4-S246A-R | TCGGTTCAGCAGCTGTTTTCCTAAGAGGGAAGTCA |
| HDAC4-S467A-F | ACCCAGGCGGCCCCGCTGCCCCAGAACGCCCA |
| HDAC4-S467A-R | AGCGGGGCCGCCTGGGTCCGCCCCAGTGGGCG |
| HDAC4-S632A-F | AGTCCGCACCCGCGTCTGCCACCTTCCCCGTG |
| HDAC4-S632A-R | AGACGCGGGTGCGGACTGCGCCCGGGACAGAG |
| N-HA-PP1α-F | GGAGGCCCGAATTCGGTCGACAATGTCCGACAGCGA GAAGC |
| N-HA-PP1α-R | CATGTCTGGATCCCCGCGGCCGCTCATTTCTTGGCTT |

| | |
|---|---|
| | TGGCGG |
| N-HA-PP1β-F | GGAGGCCCGAATTCGGTCGACAATGGCGGACGGGG AGCT |
| N-HA-PP1β-R | CATGTCTGGATCCCCGCGGCCGCTCATCACCTTTTCT TCGGCG |
| N-HA-PP1γ-F | GGAGGCCCGAATTCGGTCGACAATGGCGGATTTAGA TAAACTCAAC |
| N-HA-PP1γ-R | CATGTCTGGATCCCCGCGGCCGCTCATTTCTTTGCTT GCTTTGTGA |
| GSDMD-K103R-F | ACAGGCAAGGATCGCAGGCGGGGCCGCGGTGT |
| GSDMD-K103R-R | CTGCGATCCTTGCCTGTCCTGGGGCTGCCAGC |
| GSDMD-K145R-F | AGAACACAGAGTCCTGCAGCAGCTGCGCAGCC |
| GSDMD-K145R-R | GCAGGACTCTGTGTTCTGGCTGCCGCAGGTGC |
| GSDMD-K248R-F | AGGCCACAGGCGTTCCACGAGCGAAGGCGCCT |
| GSDMD-K248R-R | TGGAACGCCTGTGGCCTGTCGCGGGTGGCTGG |
| GSDMD-K387R-F | AAACGCAGCACAGGCTGCTGGCGGAGGCGCTGGAG T |
| GSDMD-K387R-R | CAGCCTGTGCTGCGTTTCACTCAGCATGGTCA |
| PLVX-IRES-h-GS DMD-FL-F | GGATCTATTTCCGGTGAATTCATGGGGTCGGCCTTTG AG |
| PLVX-IRES-h-GS DMD-FL-R | CTTGTAGTCGGATCCGCGGCCGCTGTGGGGCTCCTG GCTCA |

**Supplementary Table 2. siRNA sequences for the mouse HDAC4 oligonucleotide.**

| Name | Sense (5’-3’) |
|---|---|
| si-HDAC4-#1-F | CACCAUCCUUACCCAACAUTT |
| si-HDAC4-#1-R | AUGUUGGGUAAGGAUGGUGTT |
| si-HDAC4-#2-F | GCGACACCAUAUGGAAUGATT |
| si-HDAC4-#2-R | UCAUUCCAUAUGGUGUCGCTT |
| si-HDAC4-#3-F | GGCUGAAUGUGAGCAAGAUTT |
| si-HDAC4-#3-R | AUCUUGCUCACAUUCAGCCTT |
